# TrACES of Time: Towards estimating time-of-day of bloodstain deposition by targeted RNA sequencing

**DOI:** 10.1101/2025.02.03.636230

**Authors:** Annica Gosch, Sebastian Sendel, Amke Caliebe, Cornelius Courts

**Affiliations:** Institute of Legal Medicine, University Hospital Cologne, Cologne, Germany; Institute of Medical Informatics and Statistics, Kiel University and University-Hospital Schleswig-Holstein, Kiel, Germany

**Keywords:** Forensic RNA Analysis, trace contextualisation, targeted RNA sequencing, time-of-day, gene expression

## Abstract

**Background:** In forensic molecular biology, the main task consists of identifying individuals who contributed to biological traces recovered from (potential) crime scenes. However, to support evidence-based reconstruction of the course of activities having taken place at the scene, contextualising information regarding how and when a biological trace was deposited is oftentimes required.

**Results:** Here we present the development of a forensic molecular biological analysis procedure for the prediction of the time-of-day at which a bloodstain has been deposited by targeted quantification of selected mRNA markers. Time-of-day candidate prediction markers with diurnally rhythmic expression have previously been identified by whole transcriptome sequencing. Here, we build on our previous findings by establishing a targeted cDNA sequencing protocol on an Ion S5 massively parallel sequencing device for the targeted gene expression quantification of 74 time-of-day candidate prediction markers. Based on expression measurements of these markers in 408 blood samples (from 51 individuals deposited at eight time points over a day), we establish and compare different statistical methods to predict time of deposition. The most suitable model employing penalised regression achieved a root mean squared error of 3 hours and 44 minutes with 78 % of predictions being correct within +/- 4 h (evaluated by five-fold cross-validation).

**Conclusions:** Our study provides the first prediction model for time-of-day of bloodstain deposition based on targeted RNA sequencing and thus represents an important step towards forensic trace deposition timing. It thereby relevantly contributes to the growing knowledge on <u>Tr</u>anscriptomic <u>A</u>nalyses for the <u>C</u>ontextualisation of <u>E</u>vidential <u>S</u>tains (TrACES).

## 1. Background

The main task for forensic molecular biology is to identify individuals who contributed to biological traces recovered from (potential) crime scenes. However, to objectively reconstruct the course of events and activities having taken place at the scene, contextualising information regarding how and when a biological trace was deposited is oftentimes required but can typically not be gathered from standard forensic genetic procedures.

Due to its adaptive and dynamic nature, the transcriptome (i.e. the entirety of RNA molecules expressed in the cells contained in biological material/fluids) is frequently targeted to obtain such contextualising information [1]. Several forensic genetic laboratories routinely analyse differentially and tissue/fluid specifically expressed transcripts to assess the cellular compositions of biological traces [2, 3]. Beyond this, the potential of transcriptomic analyses to support the investigation of other aspects of forensic relevance is explored in several current research projects, e.g. focusing on transcriptomic changes after wound infliction to estimate the wound age [4, 5], changes occurring after death of an organism to estimate the post-mortem interval [6, 7] or ageing-related changes suitable for the estimation of the biological age of a trace donor [8].

A question of particular forensic relevance is the time of trace deposition. Information about this time may help to determine when a certain criminal activity has taken place but also helps to assess which traces recovered at a crime scene are actually relevant for the reconstruction of a specific event. Such insights may be helpful both in the *investigative* (i.e. to prioritise investigative leads) as well as in the *evaluative* (i.e. to assess the plausibility of how events unfolded according to the hypotheses of the prosecution and/or the defence within a criminal trial) stage [9]. Scientific studies targeting the transcriptome have approached the question of trace deposition timing in two different ways: First, by estimating the *time* elapsed *since* the trace has been deposited, for instance by measuring the integrity of one or several specific RNAs (’differential degradation’) [10, 11], and second by quantifying RNA markers with diurnally oscillating expression to estimate the *time-of-day* of stain deposition [12].

Estimating the time-of-day of stain deposition using rhythmically expressed RNA markers was first attempted by Lech et al. [12, 13]. In 2014, they analysed two different microRNA (miRNA) markers which had previously been reported to show significant expression differences in vitreous humour from deceased individuals depending on the time-of-day the individual died. They did not observe significant expression differences for these two markers between venous blood samples collected from twelve male individuals at different timepoints over the day [13]. When looking at mRNA markers in the same blood samples, however, Lech et al. observed significantly rhythmic expression for eleven out of 21 analysed markers. Based on gene expression measurements from these selected mRNA markers (alone or in combination with two hormonal biomarkers melatonin and cortisol), they built statistical models allowing for a distinction of three time-of-day categories (night/early morning, morning/noon and afternoon/evening) with AUCs between 0.75 and 0.95 [12]. Since then, two research groups have further explored the potential of rhythmic mRNA marker analysis for predicting the time-of-day of bloodstain deposition: Kirk et al. [14] assessed expression patterns of ten preselected candidate mRNA markers in dried blood spots collected from ten individuals, but observed no significant differences between time points of sample deposition, except for a single marker. The authors acknowledged the possibility of more suitable markers being identified in future approaches, possibly by whole transcriptome sequencing. Most recently, Cheng et al. [15] assessed the expression patterns of seven preselected mRNA markers in dried bloodstains from eight individuals and reported significant differences between time-of-day of sample collection for four mRNA markers. The authors devised and compared several statistical models to predict time-of-day of sample collection based on expression measurements of these four mRNA markers and achieved an RMSE of 5.98 h and a mean absolute prediction error of 3.92 hours for their best performing model using the k-nearest neighbour algorithm. In these three studies, partly overlapping sets of 21, ten and seven candidate mRNA markers (constituting 24 different mRNA markers) were evaluated. Given the low number of markers assessed in each study (limited by the use of quantitative PCR (qPCR) for gene expression quantification) and a potential bias of the marker selection processes (markers selected from previously published research with study conditions mostly not reflecting realistic, forensic casework-like settings), we deemed it possible that some potentially well-performing time-of-day prediction candidate markers might not have been investigated yet. In our research project “TrACES” (<u>Tr</u>anscriptomic <u>A</u>nalyses for the <u>C</u>ontextualisation of <u>E</u>vidential <u>S</u>tains) [16], we thus intended to approach these limitations by identifying suitable time-of-day prediction markers in a completely independent approach sequencing the entire transcriptome of blood samples deposited at different timepoints of a day. Based on a dataset of 80 samples from ten different individuals spending their day in an uncontrolled (i.e. forensically relevant) manner, we identified 81 RNA markers exhibiting diurnally rhythmic expression oscillations.

In the present study, building on our previous findings, we assessed the performance of these newly identified candidates in the prediction of bloodstain deposition time. The development of a targeted RNA sequencing approach applicable to bloodstains commonly encountered in forensic settings enabled us to simultaneously quantify the expression of 69 candidate predictors in a well-curated sample set of 408 dried blood samples collected from 51 individuals at eight different time points of the day. The markers with significantly rhythmical expression were then used to construct and compare different statistical models for predicting the time-of-day of blood stain deposition and different factors impacting model performance were evaluated.

## 2. Results

In the present study, we analysed a total of 408 blood samples obtained from 51 healthy individuals deposited at eight different time points of the day (8 AM, 11 AM, 2 PM, 5 PM, 8 PM, 11 PM, 2 AM and 5 AM). In each sample, gene expression of candidate time-of-day prediction markers was quantified by targeted RNA sequencing and expression measurements were normalised with two alternative approaches (based on total reads in a sample (CPM-normalisation) and based on reference genes (Refall-normalisation)).

Originally, a total of 84 time-of-day candidate prediction markers were analysed in this study. Initial evaluation of marker suitability considering expression level, amplification performance, significance of expression differences between timepoints as well as correlation between markers led to the exclusion of a total of 15 candidate markers from each normalised dataset (cf. 5.2 and 5.3), resulting in two datasets with normalised expression data for 69 time-of-day candidate prediction markers, each.

### 2.1 Development and evaluation of statical models for time-of-day prediction

We developed prediction models for time-of-day of sample deposition based on gene expression measurements by four different statistical approaches, including machine learning methods (Penalised Regression (PR), Random Forest (RF), Support Vector Machine (SVM) and a previously published approach named TimeSignatR [17]). Time was either modelled as a categorical (dividing a day into eight categories) or as a continuous variable (cf. 5.3.3). To be able to apply the prediction models to the continuous and cyclic variable time-of-day, time was treated as the angle of the hour hand of a 24-hour clock and subsequently trigonometrically transformed to a sine and a cosine component, which were then modelled separately (Figure 1).

**Figure 1:**
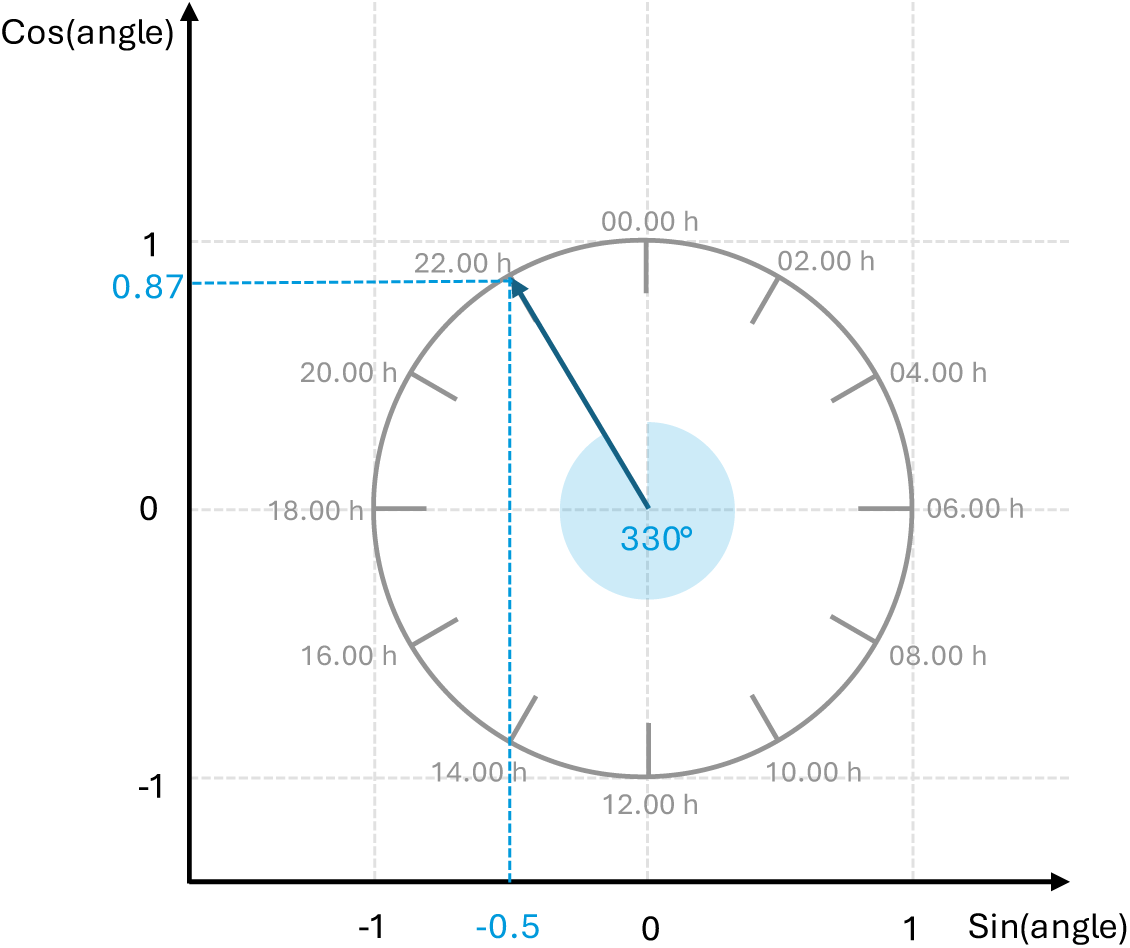
Visualisation of the trigonometric transformation of time-of-day for continuous modelling. Time is considered as the angle of the hour hand on a 24-hour-clock (represented by a unit circle, shown in grey). The sine and the cosine component of the time-of-day are plotted on the x- and the y-axis, respectively. In the example depicted in this plot (shown in blue), the time is set to 22.00 h, represented by an angle of 330° on a 24-hour-clock (blue area). The arrowhead corresponds to a value of -0.5 on the x-axis (= sin(330°)) and 0.87 on the y-axis (= cos(330°)).

Table 1 shows the root mean squared error (RMSE) as estimated by five-fold cross-validation for six different statistical approaches (for both the CPM- and the Refall-normalised dataset, modelling time as either categorical or continuous). Additional performance metrics are presented in Suppl. Table 1. All models performed significantly better than a random chance predictor.

**Table 1:** Comparison of model performance (RMSE) for six modelling approaches in each normalised dataset.

| Normalised dataset |  |  | CPM |  |  |  |  |  | Refall |  |  |  |  |  |  |
| --- | --- | --- | --- | --- | --- | --- | --- | --- | --- | --- | --- | --- | --- | --- | --- |
| Time modelled |  |  | continuously |  |  |  | categorically |  | continuously |  |  |  | categorically |  |  |
| Prediction model |  |  | TS | PR | RF | SVM | RF | SVM | TS | PR | RF | SVM | RF | SVM | RP |
| No. of samples |  |  |  |  |  |  |  |  |  |  |  |  |  |  |  |
| RMSE/h | Overall | 408 | 3.76 | 3.79 | 4.05 | **3.73** | 4.66 | 4.03 | **3.71** | 3.73 | 3.91 | **3.71** | 4.59 | 4.13 | 6.93 |
|  | 02.00h | 51 | **3.34** | 3.42 | 4.02 | 3.43 | 4.70 | 4.05 | **3.22** | 3.28 | 4.06 | 3.60 | 4.30 | 4.24 | 6.93 |
|  | 05.00h | 51 | **3.06** | 3.16 | 4.48 | 3.84 | 4.52 | 3.92 | **3.13** | 3.26 | 4.25 | 3.44 | 4.60 | 3.76 | 6.93 |
|  | 08.00h | 51 | 2.32 | 2.36 | 2.54 | **1.95** | 2.66 | 2.45 | 2.47 | 2.55 | 2.52 | 1.87 | 3.17 | **1.68** | 6.93 |
|  | 11.00h | 51 | 3.15 | 3.20 | 3.65 | **3.02** | 4.37 | 4.01 | 3.20 | 3.33 | 3.68 | 3.01 | 4.49 | **3.00** | 6.93 |
|  | 14.00h | 51 | 4.01 | 4.21 | 3.77 | **3.74** | 4.43 | 4.09 | 4.18 | 4.09 | **3.45** | 3.53 | 4.56 | 4.35 | 6.93 |
|  | 17.00h | 51 | 4.54 | 4.61 | 4.87 | **4.44** | 5.08 | 4.79 | 4.57 | 4.35 | **4.16** | 4.44 | 5.04 | 5.31 | 6.93 |
|  | 20.00h | 51 | 4.54 | **4.27** | 4.47 | 4.43 | 5.64 | 4.37 | **4.13** | 4.18 | 4.58 | 4.56 | 5.25 | 5.04 | 6.93 |
|  | 23.00h | 51 | 4.49 | 4.51 | 4.15 | 4.33 | 5.30 | **4.20** | 4.26 | 4.36 | **4.18** | 4.47 | 4.97 | 4.45 | 6.93 |

The RMSE estimated by five-fold cross-validation for four different statistical models (Penalised Regression (PR), Random Forest (RF), Support Vector Machine (SVM) and TimeSignatR (TS)) is presented for the entire dataset (n=408 samples from 51 individuals deposited at eight different timepoints, dried for 48 hours) for all time points (overall) as well as for samples collected at each timepoint separately (51 samples per timepoint). Raw gene expression measurements were normalised to total read counts per sample (CPM) or the expression of selected reference genes (Refall). The best result per normalised dataset is indicated in bold. The last column indicates prediction performance achieved if a random predictor (RP) was used.

In a direct comparison between the categorical models and their continuous counterparts (SVM and RF), both categorical classifiers resulted in higher RMSE compared to the continuous model (Table 1). Therefore, the continuous modelling approach was considered more suitable.

When looking at the individual predictions for the four different continuous models, similar results were obtained for the PR model and the TimeSignatR algorithm, which was expectable as TimeSignatR is also based on penalised regression. In both datasets, individual predictions obtained by the PR/TimeSignatR model were more similar to the SVM results than results obtained by the RF algorithm (Suppl. Figure 1).

For both normalisation procedures, the best overall performance was observed for SVM and PR/TimeSignatR. A closer analysis of the performance metrics per sample deposition time point (Table 1, Suppl. Table 1) showed that Penalised Regression and TimeSignatR produced the most consistent results across all time points, whereas the greatest differences were observed for the RF models.

Due to this greater consistency and the better general comprehensibility and explainability of PR compared to SVMs, PR was determined to be the overall most suitable model (with TimeSignatR and PR being considered equivalent, PR was chosen as the more general approach).

In accordance with expectations based on preliminary testing (Suppl. File 1), similar results were obtained for both normalisation procedures (Table 1, Suppl. Table 1). Refall-normalisation can be achieved with quantification values from all reference genes and the target gene of interest and normalised values are hence independent of all other markers in the same sequencing panel whereas CPM normalisation requires calculation of the total read count for all markers in the same panel. Therefore, all further evaluations were solely based on the Refall-normalised dataset. Individual predictions for all 408 samples after five-fold cross-validation for the final PR model are presented in Figure 2a.

**Figure 2:**
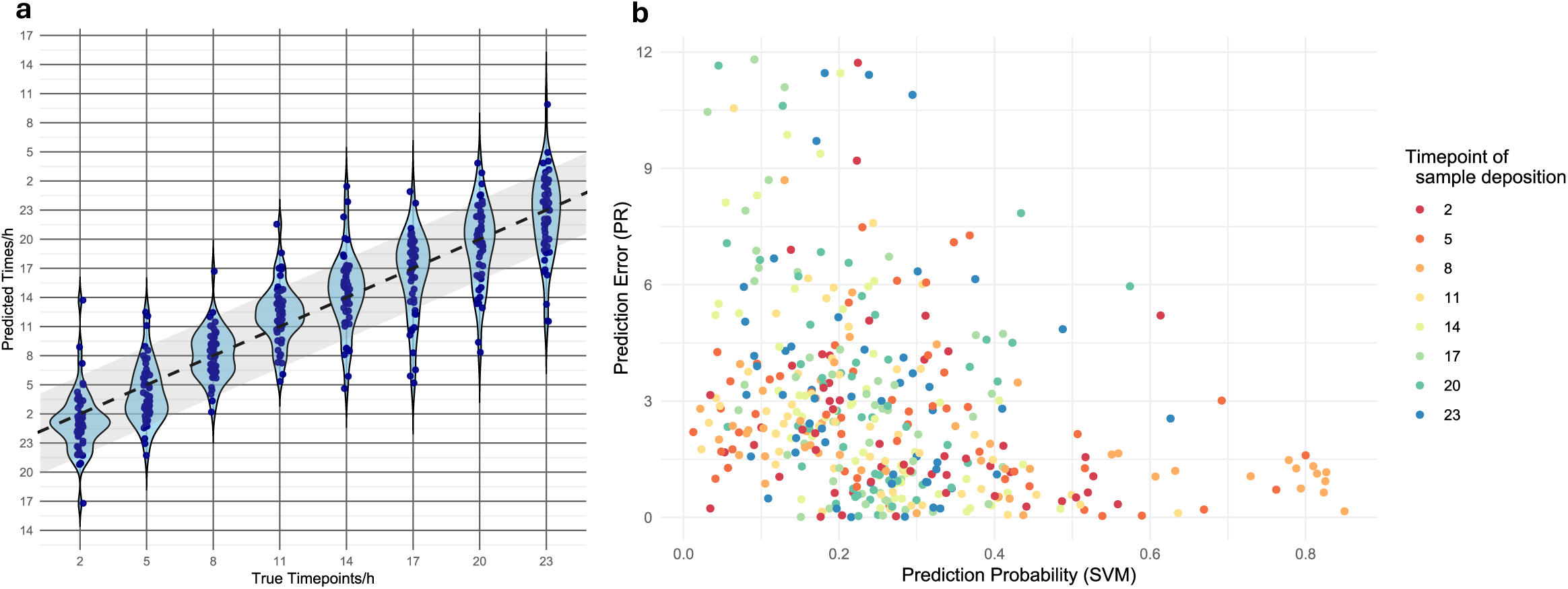
Predictions obtained from the final model (PR) **a** – Violin plot showing the individual predictions (dark blue) and the kernel density plots (light blue) obtained for the individual samples (n = 408 samples from 51 individuals deposited at eight different timepoints, dried for 48 hours) in the respective test set of five-fold cross-validation with the final PR model (Refall-normalised data). The area of predictions correct within +/- 4 hours is shaded in light grey and x = y-values are shown as a dashed line in black. Note that in order to be able to depict full violin plots per sample deposition time point, the y-axis was stretched over more than one day.**b** – Scatterplot showing the relationship between the absolute prediction errors (obtained for the final continuous PR model (on the Refall-normalised dataset)) and the probabilities for the corresponding category (obtained from the categorical SVM model (Refall-normalised dataset)) for n = 408 samples from 51 individuals deposited at eight different timepoints, dried for 48 hours. Colours indicate the true sample deposition time point.

In contrast to continuous regression models, the output of a categorical classifier contains prediction probabilities for each predicted category as a measure of uncertainty of the individual prediction. Hence, in a next step, we aimed to evaluate whether probabilities obtained for the best performing categorical model (SVM) can be used to obtain information about the reliability of individual predictions obtained from the continuous PR model. Figure 2b shows that for very high categorical probabilities (> 0.7), prediction errors were consistently low (< 2 hours) and the highest prediction errors (> 10 hours) were associated with low categorical prediction probabilities (< 0.3). However, there was also a large proportion of samples with low prediction errors despite showing a low probability. Overall, the prediction error and categorical prediction probability were significantly correlated with a Spearman’s rank correlation rho of -0.396 (pSpearman < 2,2×10^-16^). Additional but limited informative value may thus be gained from considering the distribution of categorical prediction probabilities (estimated from the categorical SVM model) in combination with the predicted timepoint (obtained from the continuous PR model). Continuous predictions obtained from the PR model in combination with categorical probabilities as estimated by the categorical SVM model for each individual sample are presented in Supp. Table 2.

**Table 2:** Coefficients for each marker in the final most suitable model (PR)

|  | Model component |  |
| --- | --- | --- |
|  | Sinus | Cosinus |
| ABCD2 | 0.000 | -0.191 |
| ABHD5 | 0.000 | 0.000 |
| ALOX5AP | -0.005 | 0.000 |
| APMAP | -0.129 | 0.161 |
| ARG1 | 0.962 | -0.280 |
| AVIL | -4.415 | 0.000 |
| BACH2 | 0.000 | 0.000 |
| BCL2 | 0.267 | 0.487 |
| CAMKK1 | 1.406 | -1.551 |
| CHST15 | 0.000 | 0.000 |
| CLEC4E | 0.063 | 0.090 |
| CLIC3 | -0.402 | -1.185 |
| CNTLN | -0.165 | -0.193 |
| CPD | -0.049 | 0.000 |
| CRISPLD2 | 0.084 | 0.000 |
| DAAM2 | 0.000 | -1.506 |
| ECHDC3 | 0.000 | 0.029 |
| ENHO | 0.000 | 12.293 |
| ERGIC1 | 0.000 | -0.246 |
| ERMN | 0.000 | 0.211 |
| FAAH2 | 0.914 | 2.043 |
| FBXL16 | 1.425 | 2.792 |
| FKBP5 | 0.250 | -0.045 |
| FLT3 | 0.000 | 0.000 |
| FOSL2 | -0.012 | 0.000 |
| GABARAPL1 | 0.000 | 0.000 |
| GCNT4 | -0.665 | 0.000 |
| HAL | -0.179 | 0.000 |
| HDAC9 | -0.541 | 0.294 |
| HNRNPDL | 0.000 | 0.000 |
| HPCAL4 | 0.000 | 0.936 |
| ID3 | -0.779 | -0.114 |
| IGF2R | -0.105 | 0.000 |
| IL13RA1 | 0.847 | 0.000 |
| IL18R1 | 0.000 | 0.000 |
|  | Sinus | Cosinus |
| IL1R1 | -0.710 | 0.000 |
| IL1R2 | 0.000 | 0.000 |
| KLF9 | 0.158 | -0.293 |
| LEF1 | 0.000 | 0.000 |
| MAL | 0.029 | 0.084 |
| MFGE8 | 0.000 | 0.000 |
| MGAM | -0.068 | 0.000 |
| MKNK2 | 0.000 | 0.000 |
| MSRB2 | -0.021 | 0.000 |
| MTHFS | 0.000 | 0.000 |
| MYO1E | 1.066 | 2.749 |
| NELL2 | -0.015 | 0.000 |
| NR1D1 | -0.361 | 0.017 |
| NR1D2 | 0.000 | 0.079 |
| NRCAM | 1.266 | 0.000 |
| PER1 | 0.000 | 0.101 |
| PER2 | 4.937 | 0.000 |
| PER3 | 2.763 | -0.463 |
| PFKFB2 | -0.048 | 0.297 |
| PHC2 | 0.000 | 0.000 |
| SAP30 | 0.000 | 0.000 |
| SEC14L1 | 0.012 | 0.051 |
| SIPA1L2 | 0.000 | 0.000 |
| SMAP2 | 0.069 | 0.000 |
| SPATA6 | 0.168 | 0.000 |
| SYTL3 | -0.085 | 0.000 |
| TCF7 | 0.000 | 0.224 |
| TMCC3 | 0.000 | -0.465 |
| TPST1 | 0.000 | -1.239 |
| TRABD2A | -0.416 | -0.469 |
| TSC22D3 | 0.011 | -0.049 |
| UPB1 | 4.083 | -2.867 |
| XKR8 | 0.000 | 0.551 |
| ZBTB16 | -0.888 | -0.299 |

Notably, it was observed that all algorithms provided the most accurate results for samples deposited at 8 AM and generally more accurate results for samples deposited late at night or in the morning, whereas predictions for samples deposited in the afternoon/evening were less accurate (Table 1, Figure 2a).

Model coefficients assigned to the 69 candidate time-of-day prediction markers in the final most suitable prediction model (PR) are shown in Table 2. Higher absolute values indicate a higher importance of the respective marker in the model. The sine component of the model was composed of 28 and the cosine component of 33 markers. Several markers were relevant only in the sine or the cosine component of the model, whereas only 15 markers were not included in either model component.

Coefficients were extracted for a PR model trained on the entire Refall-normalised dataset (n = 408 samples from 51 individuals deposited at eight different timepoints, dried for 48 hours). Markers excluded from the respective model (coefficient = 0) are highlighted in grey, the top ten most relevant markers in each model are highlighted in orange.

### 2.2 Model evaluation: donor properties

In a previous study analysing whole transcriptome sequencing data [16], we observed that time-of-day prediction accuracies were improved when accounting for interindividual differences in gene expression by normalising expression measurements across all samples from one individual (as implemented in the original TimeSignatR algorithm [17]). A similar improvement was observed in this study: When training the PR model on gene expression measurements renormalised within each subject (after Refall-normalisation), we obtained an RMSE of 2.62 h estimated by five-fold cross-validation, representing a reduction of RMSE by approximately 67 minutes compared to the model without within-subject renormalisation (Figure 3). This demonstrates that prediction accuracies are relevantly impacted by interindividual differences.

**Figure 3:**
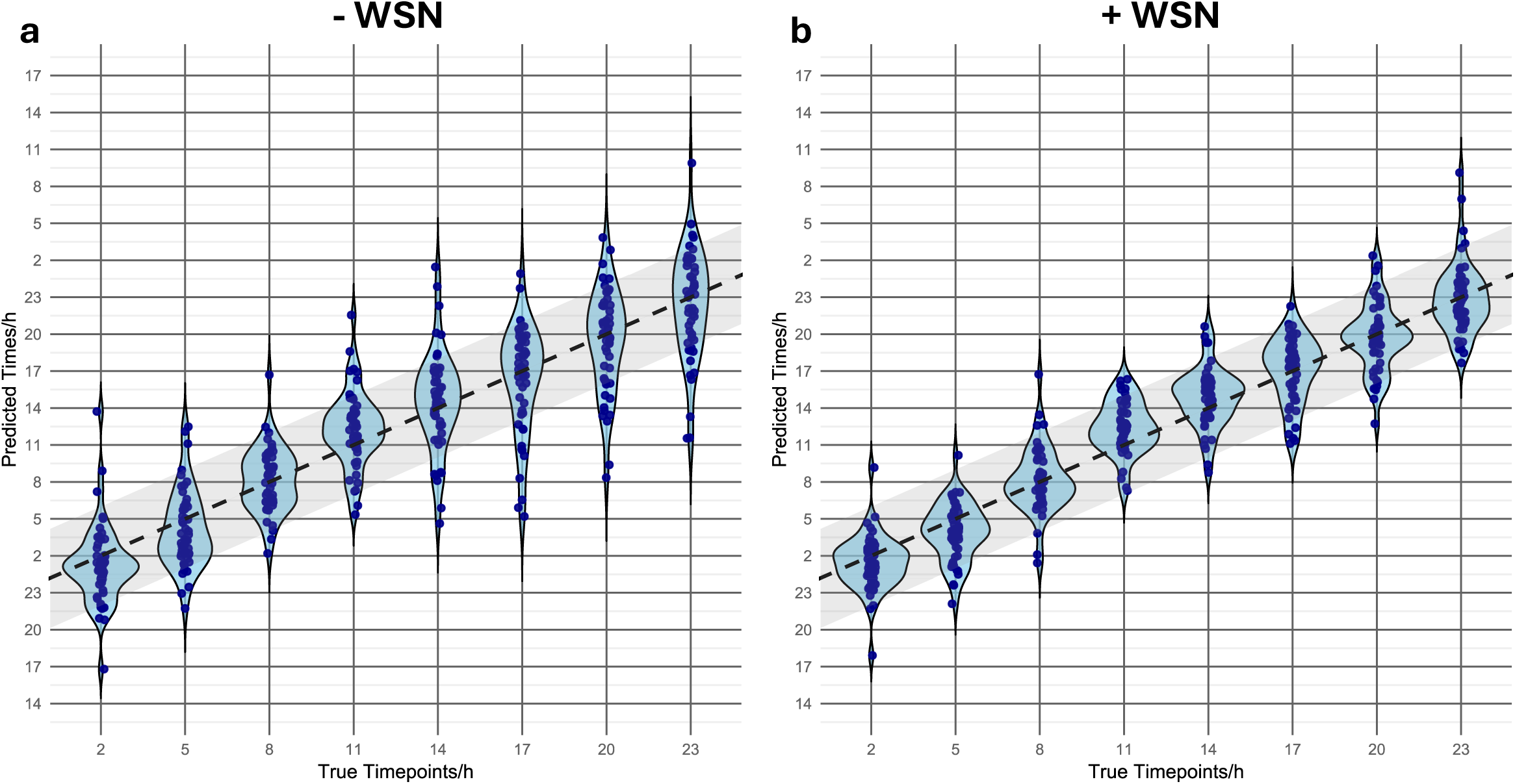
Violin plots for time-of-day predictions without (-WSN) and with within-subject renormalisation (+WSN) **a** – Violin plot showing the individual predictions (dark blue) and the kernel density plots (light blue) obtained for the individual samples (n = 408 samples from 51 individuals deposited at eight different timepoints, dried for 48 hours) in the respective test set using five-fold cross-validation with the final PR model (Refall-normalised data without WSN). The area of predictions correct within +/- 4 hours is shaded in light grey and x=y-values are shown as a dashed line in black. **b** – Violin plot showing the individual predictions (dark blue) and the kernel density plots (light blue) obtained for the individual samples (n = 408 samples from 51 individuals deposited at eight different timepoints, dried for 48 hours) in the respective test set using five-fold cross-validation with a PR model (Refall-normalised data with WSN). The area of predictions correct within +/- 4 hours is shaded in light grey and x=y-values are shown as a dashed line in black. Note that in order to be able to depict full violin plots per sample deposition time point, the y-axis was stretched over more than one day.

Certain information about stain donor characteristics, such as age and sex, may be obtained by routinely performed forensic molecular biological analysis methods. Therefore, we tested whether including this information in our prediction model would increase model performance. However, including donor age and sex information did not lead to a relevant increase in prediction accuracy: Using five-fold cross-validation, we obtained an RMSE of 3 hours and 39 minutes (representing a decrease by five minutes compared to the model not considering donor information). In accordance with these results, we did not observe significant differences between prediction errors obtained for male and female or differently aged individuals (pLMM > 0.05 for both age and sex).

Additionally, we assessed the impact on prediction accuracy of other donor characteristics and activities on which information was available for participants in this study. As this study was not set-up to systematically investigate the impact of these donor characteristics and donor activities on prediction accuracy, most characteristics were represented only a few times in our dataset. Thus, the investigations described below have to be considered exploratory in nature: Suppl. Figure 2 represents the observed prediction errors for our final PR model in relation to donor age, sex, chronotype and time of the year of sample deposition as well as individual intake of caffeine or medication and napping behaviour prior to sample deposition. In accordance with previous observations, no differences in prediction error were observed between male and female participants (Suppl. Figure 2a) or between participants from different age groups (Suppl. Figure 2b). No differences in prediction error became apparent for individuals with different chronotypes (Suppl. Figure 2c). Visual inspection seemed to suggest slightly lower prediction errors for samples collected in spring as compared to winter and autumn (Suppl. Figure 2d), however, both factors did not significantly impact prediction error (pLMM >0.5). Several individuals stated having consumed caffeine within the last three hours prior to sample deposition. Prediction errors did not seem to differ between samples with and without prior caffeine consumption (Suppl. Figure 2e, pLMM >0.5). Study participants were instructed to refrain from consuming alcohol on the day of study participation. A single study participant notwithstanding reported having consumed a single glass of wine in the evening. A low prediction error of 1.1 hours was computed for the sample deposited after alcohol consumption (Suppl. Figure 2f), but this single observation does not warrant any conclusions about the general impact of alcohol consumption. Additionally, despite being instructed to stay awake for the day of study participation, seven participants reported having taken naps prior to depositing the 2AM or 5AM samples. Prediction errors did not seem to be increased for samples collected after napping (Suppl. Figure 2f), yet again due to the low number of occurrences, no general conclusions about the impact of napping can be drawn. Lastly, comparable prediction errors were observed for individuals taking certain medication (Suppl. Figure 2g). As before, the variability of medication taken by study participants in combination with the low number of participants taking each individual medication precludes valid general conclusions.

In a last approach, we assessed how well true and predicted timepoints of sample deposition aligned for each individual over the entire day. In this approach, we noted that individuals could be broadly divided into three groups: individuals for which true and predicted timepoints generally aligned well, individuals for which there are larger deviations only for a certain period of the day and individuals for which true and predicted time generally did not align well. Plots showing true and predicted times for each of the three groups (hereafter named “consistent match group”, “intermittent match group” and “mismatch group”) are shown in Suppl. Figure 3a-c.

A rational assumption would be that prediction errors in the “mismatch group” most likely arise from general donor characteristics whereas prediction errors in the “intermittent match group” are more likely to have arisen from individual activities. We thus explored whether donors with certain characteristics or having performed certain activities were more likely to be categorised in the “intermittent match group” or the “mismatch group”. As most observations were consistent with previous descriptions, we report only a selection of observations that we consider worth noticing: Both study participants > 50 years of age were not in the “consistent match group”, however, only one of them was categorised into the “mismatch group”. Only one of the participants with extreme chronotypes (definite morning (n = 4) or definite evening (n = 1)) was included in the “mismatch group” (the others were part of the “consistent match group” (n = 2) or the “intermittent match group” (n = 2)). All nine participants who reported the intake of medication on the day of study participation (excluding vitamins and oral contraceptives) were either in the “mismatch” or the “intermittent match group”, yet the time periods for which the largest prediction errors were observed did not necessarily correspond to the time of medication intake (Suppl. Table 3). Notably, for the individual with the highest average prediction error (v027: mean absolute error of 5.4 hours) no exceptional individual characteristic possibly explaining these deviations could be identified (male individual, 29 years, intermediate chronotype, no medication, no extraordinary food consumption or physical activity on the day of sample deposition).

### 2.3 Model evaluation: sample properties

#### 2.3.1 Aged samples

The impact of storage duration was assessed by directly comparing predictions obtained for sample aliquots obtained from the same individual at the same deposition time point and stored for either 48 hours or 16 days (prior to being kept at -80 °C until further processing). A strong correlation was observed between predictions obtained for the 48 h and 16 d old samples, for both the sine (Pearson’s correlation coefficient of 0.90, pPearson < 2,2×10^-16^) and the cosine (Pearson’s correlation coefficient of 0.92, pPearson < 2,2×10^-16^) component of time-of-day. Differences in predicted times were < 3 hours for 87 % of samples (Figure 4). For samples stored for 48 h, an RMSE of 4.07 h was obtained. Samples stored for 16 d resulted in an RMSE of 4.33 h, indicating an increase by approximately 16 minutes.

**Figure 4:**
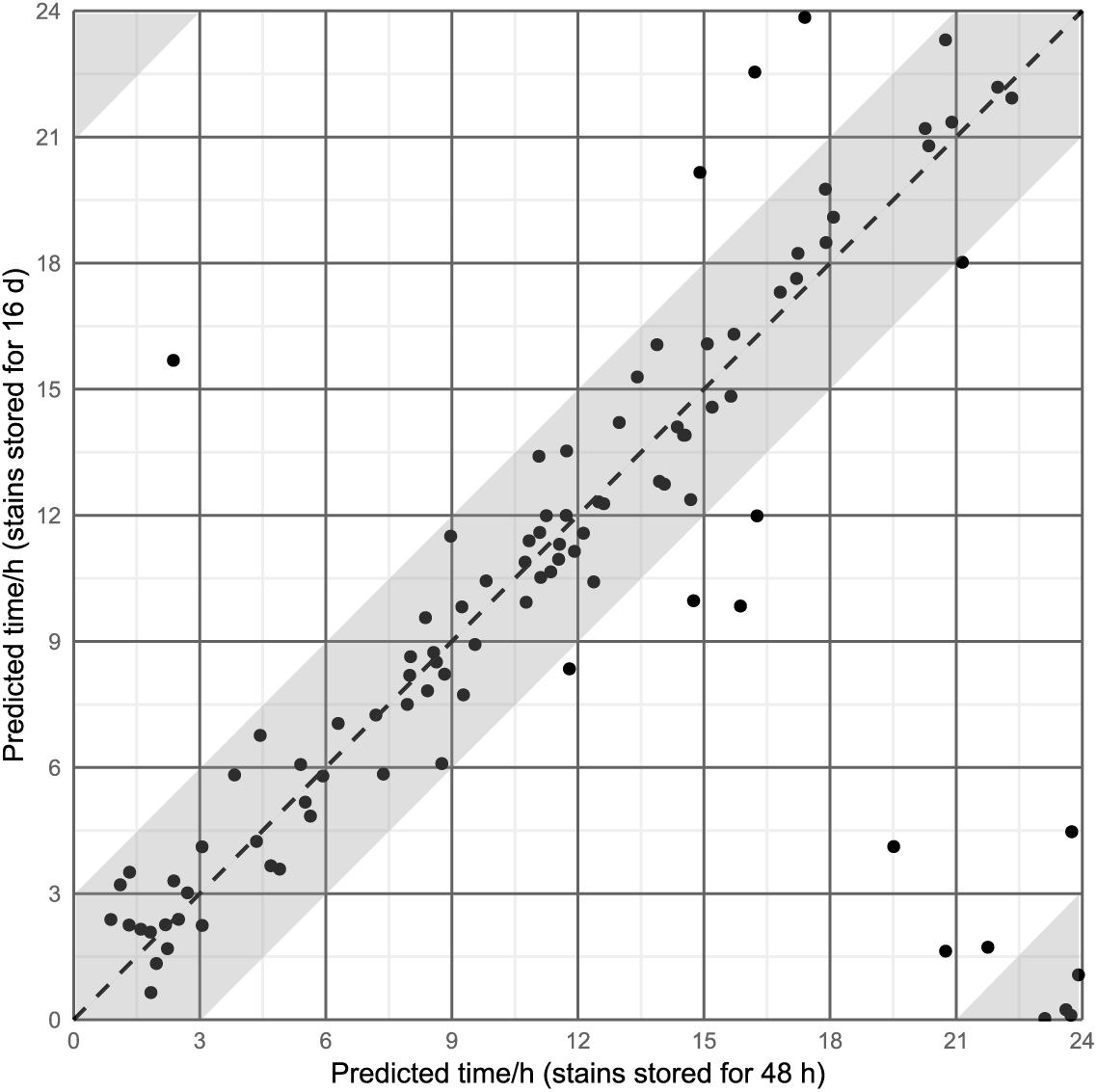
Predictions obtained for 48 h and 16 day old samples. Scatterplot showing the relationship between predictions obtained for stains stored for 48 hours (x-axis) and 16 days (y-axis) for n = 100 samples obtained from 25 individuals deposited at four different timepoints. The area of predictions close to each other within +/- 3 hours is shaded in light grey and x = y-values are shown as a dashed line in black.

#### 2.3.2 Diluted samples

The impact of sample dilution was assessed by directly comparing predictions obtained for sequencing libraries prepared from RNA extracts with different RNA input amounts. A significant correlation was observed between predictions obtained for input amounts of 25 and 5 ng, for both the sine (Pearson’s correlation coefficient of 0.73, pPearson = 0.007) and the cosine (Pearson’s correlation coefficient of 0.92, pPearson = 0.00002) component of time-of-day, while the correlation between input amounts of 25 and

0.5 ng RNA was not significant, for neither the sine (Pearson’s correlation coefficient of 0.55, pPearson = 0.06) nor the cosine (Pearson’s correlation coefficient of 0.49, pPearson = 0.1) component of time-of-day. Predicted times for diluted and undiluted samples are shown in Figure 5. The RMSE increased with sample dilution, ranging from 3.58 h for samples with 25 ng RNA input, 3.74 h for samples with 5 ng RNA input, to 4.55 h for samples with 0.5 ng RNA input.

**Figure 5:**
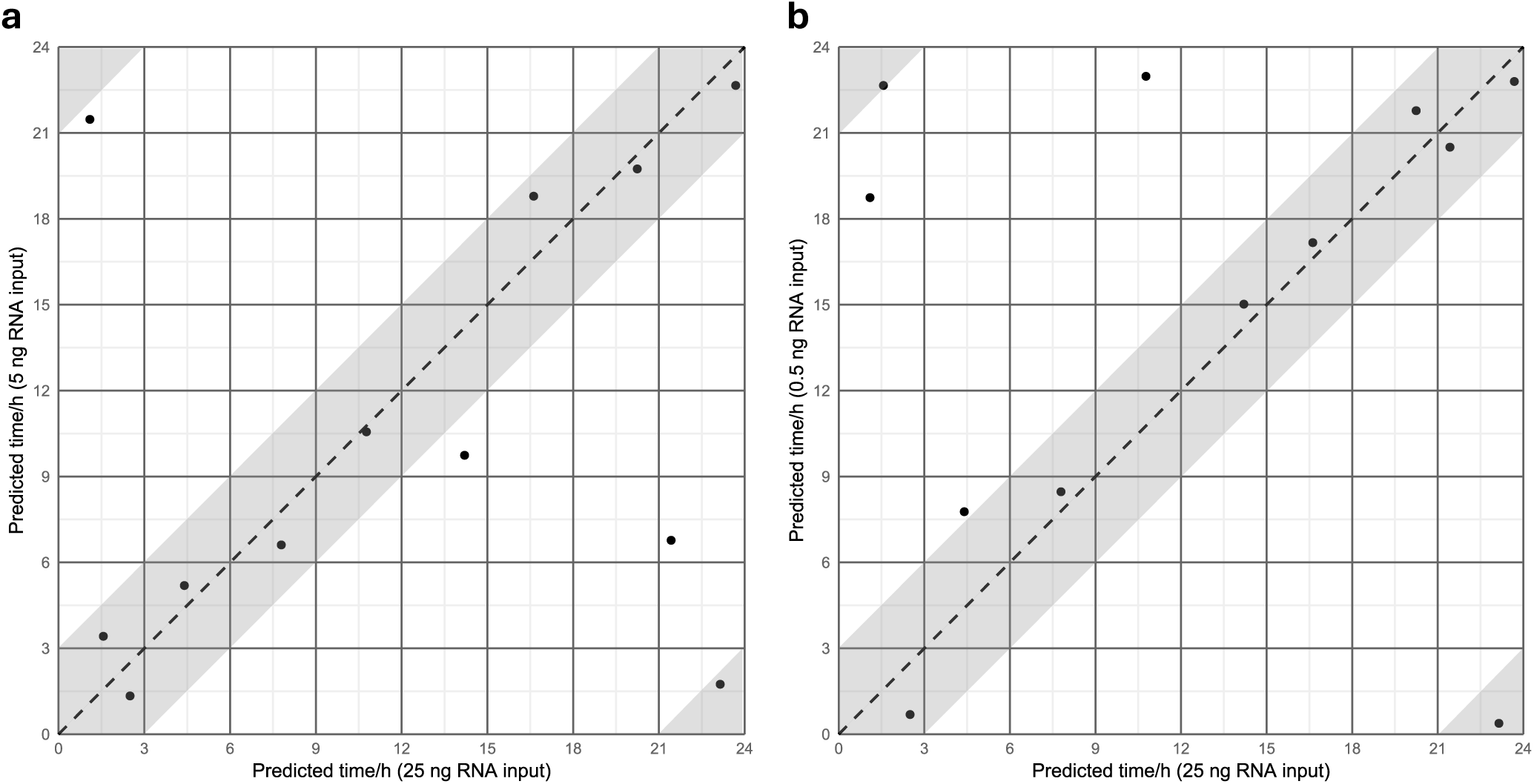
Predictions obtained for samples with optimal and sub optimal R A input amounts. Scatterplot showing the relations ip between predictions obtained for stains sequenced with optimal sample input amounts 25 ng and suboptimal input amounts **a:** 5 ng, **b:** 5 ng The area of predictions close to each other within +/- 3 ours is shaded in light grey and x = y-values are shown as a dashed line in black

### 2.4 Body fluid identification (BFI)

To enable the identification of body fluid(s) present in the analysed crime scene stain, primers for the amplification of BFI markers were included in the primer panel. Suppl. Figure 4 shows the Refall-normalised expression of all 23 BFI markers in our sample set of 408 blood samples collected from 51 individuals and dried for 48 hours. In all 408 samples, all three blood markers (*ANK1, CD3G* and *SPTB*) were detected. On average, 99.6 % (range: 99.0-100 %) of Refall-normalised BFI marker reads were mapped to one of the three blood markers. As described by Hanson et al. [18], the low percentage of reads mapping to non-blood markers may be attributable to “system noise” or low-level co-expression of these markers and thus does not indicate the presence of a body fluid other than blood.

## 3. Discussion

### 3.1 Targeted sequencing for gene expression quantification

In this work, we describe the development of a targeted cDNA sequencing assay to quantify the expression of time-of-day candidate prediction markers. Targeted sequencing has been shown to facilitate multiplex analysis of over 400 markers in a single assay [19], which represents a decisive improvement compared to alternative methods for gene expression quantification such as qPCR, and has greatly advanced research in forensic body fluid identification [20–22]. As differences in gene expression between different forensically relevant body fluids/organ tissues are relatively large, the expression quantification does not need to be overly accurate [23, 24] and hence no extensive approaches to assess the gene expression quantification accuracy of targeted sequencing assays have been reported in the forensic literature so far. However, other currently investigated trace contextualisation aspects (such as time-of-day of trace deposition) are associated with more subtle differences in gene expression and thus require more accurate quantification methods. In a previously reported method assessment experiment, we observed that inter-as well as intraindividual differences in gene expression in blood samples from different deposition time points can more accurately be resolved when Unique Molecular Identifiers (UMIs) are included in the library preparation process [25]. While this newly established analysis technique proved to provide sufficiently accurate results for the purpose of our study, we did observe a clear lack of standards for performing and reporting such analyses resulting in little concordance among previous studies: While some studies describing the use of QIAGEN’s UMI-based targeted RNA sequencing protocol employed normalisation to “housekeeping genes” [26, 27] (as recommended by the manufacturer [28]), others used normalisation procedures established for whole transcriptome sequencing data [29] or did not report this information at all [30]. Information on sequencing depth has, to the best of our knowledge, never been provided.

In this study, and given the forensic scope of our approach, we have thus undertaken great effort to establish procedures ensuring the generation of robust and reproducible gene expression quantification results by optimising the sequencing depth and evaluating different read count normalisation procedures (Suppl. Files 1 and 2). Nonetheless, the cross-platform reproducibility of absolute quantification results of our analysis procedure remains to be evaluated in future work. As has previously been observed in the context of forensic DNA methylation measurements [31, 32], an adaptation of the prediction model to the measurement platform might be necessary.

### 3.2 Statistical model performance

Our final most suitable model based on penalised regression produced predictions with an RMSE of 3 hours and 44 minutes (and a mean absolute error of 2 hours and 49 minutes with 78 % of predictions being correct to within +/- 4 h, evaluated by five-fold cross-validation).

Previous studies have evaluated the possibility of predicting time of deposition in forensic blood stains based on RNA expression patterns [12, 15]. Lech et al. [12] were first to establish a prediction model based on gene expression measurements. They used a multinomial logistic regression model to obtain prediction for three broad time-of-day categories. In a sample set of 216 samples from twelve male individuals and performing leave-one-out-cross-validation (LOOCV), they obtained AUC values of 0.75-0.93 for a model including six mRNA markers and 0.88-0.95 for a model including three mRNA markers and the two hormonal markers melatonin and cortisol (quantified by radioimmunoassay analysis [33]). A direct comparison of performances of this model and our newly developed model is not straightforward, as Lech et al. used categorical models and only provided summary statistics of their cross-validation. From these, it can be deduced that 71.2 % and 76.4 % of samples were predicted correctly (i.e. assigned to the correct time-of-day category) using the mRNA and combined mRNA-hormones model, respectively. Considering the accuracy within +/- 4 hours as the most comparable performance metric (as it treats a time span of 8 hours as “correct”, corresponding to the three categories of Lech et al. [12]), a comparable but slightly higher accuracy (78 %) was obtained for the most suitable model in our study. The multinomial model (as used by Lech et al. [12]) performed worse in our study (cf. 5.3.5). However, the fact that no completely identical performance metric was obtainable, the use of liquid blood with a preserving reagent by Lech et al. [12, 34] in contrast to dried blood spots in our study, the more controlled study conditions in [12] and the reduced independence of training and test set for LOOCV (indeed, an increased accuracy of 82.4 % of predictions correct within +/- 4 hours was observed for our prediction model, when using LOOCV instead of five-fold cross-validation), hinders a direct comparability of model performances.

More recently, Cheng et al. [15] used a k-nearest-neighbour algorithm on expression measurements of four mRNA markers in a total of 64 samples (collected at 8 timepoints from different individuals) and obtained an RMSE of 5.98 h when training the model on 80 % of their data and testing it on the remaining 20 %. The RMSE were smaller for all models in our study, but due to the small number of samples in the test set (20 % of 64 samples) in [15] and questionable independence between training and test set (samples from the same individual were part of the training as well as the test set), the comparability of this performance metric is limited.

Taken together, while it is not possible to quantify the contribution of each individual methodical aspect (marker selection process, number of markers included, forensically relevant sample set, analysis method or prediction model), our study provides a relevant progression compared to previous approaches.

Notably, it was observed that all algorithms evaluated in this study provided more accurate results for samples deposited late at night or early in the morning, whereas results for samples deposited in the afternoon/evening were less accurate. Interestingly, Lech et al. had also observed the best performance metrics for their night/early morning category (23.00 h - 06.59 h) [12]. This might possibly be caused by more defined expression peaks of markers at night/early morning hours compared to other time-of-day periods and/or more heterogenous activities/behaviours of study participants in the afternoon/evening.

### 3.3 Considerations for applications to casework

Overall, the prediction accuracies obtained in this study demonstrate that time-of-day information is indeed incorporated in and can be recovered from forensic casework type blood traces. Notably, we observed pronounced inter-individual differences resulting in very accurate predictions for some individuals and markedly reduced accuracies (close to random predictions) for others. Consequently, the method in its current state cannot reliably be used to prove beyond doubt that a forensically relevant blood trace definitively has or has not been deposited at a certain time-of-day and should not be used in the evaluative stage of a criminal trial. Notwithstanding, the information that a forensically relevant blood trace is more likely to have been deposited at a certain time-of-day (while other deposition times may not be excluded but may be considered (much) less likely) may constitute valuable information in the investigative phase.

The above-described overall performance metrics summarise how the final chosen model performed for the samples collected and analysed in this study. However, when applying the developed model to an unknown sample in forensic casework, the forensic expert is more interested in getting an estimate of the uncertainty of the individual prediction at hand. Overall model performance metrics as estimated from five-fold cross-validation maybe considered a good approximation of the uncertainty associated with the prediction for an unknown sample as long as the unknown sample is sufficiently similar to the type of samples tested. This approach is commonly used for example when estimating the age of an unknown sample donor based on age-dependent methylation patterns [31, 35].

In addition to the overall model performance metrics, we found that the distribution of prediction probabilities obtained from a separately trained categorical SVM model provides supplementary information that may to a certain extent allow for an estimation of the uncertainty associated with a prediction. Notwithstanding, as categorical probabilities (especially probabilities assigned to samples collected in the afternoon/evening) often showed similarly high (and thus altogether low) values for several subsequent categories, the informative value especially of low prediction probabilities is rather limited.

Lastly, to get a good understanding of how well the developed model will perform on unknown samples e.g. encountered in forensic casework, a comprehensive understanding of additional factors influencing prediction accuracy is necessary (cf. section 3.5).

To be suitable for forensic casework, we aimed to develop a measurement method and corresponding prediction approach that can be applied to a single sample from an unknown individual, which will typically be encountered in casework scenarios. We did observe that knowing the individual, who deposited the sample, and normalising the gene expression over a set of samples collected from the same individual significantly improved prediction accuracy (Figure 3). For certain cases, especially at the time of court trial (i.e. in the evaluative stage), the person having deposited a crime-relevant blood trace is typically known. For these cases, time of deposition estimation might be improved by collecting reference samples at different timepoints and normalising gene expression over all samples obtained from the person of interest. The validity of such an approach (especially if reference samples are collected much later than the crime scene stain) would have to be evaluated in a separate study.

### 3.4 Evaluation of selected markers

An analysis of the model coefficients for each marker showed that some markers were relevant in only the sine or the cosine component of the model whereas others were relevant in both (Table 2). This observation makes sense in the context of trigonometric transformation of the variable time-of-day: Markers with expression peaks at either midnight or noon are expected to relevantly contribute to the cosine model but might show similar expression in the morning and evening and thus show model coefficients close to 0 in the sine model. Markers with expression peaks at time points lying between the four time points corresponding to maximum/minimum values in the sine and cosine model (namely 12 PM, 6 AM, 12 AM and 6 PM) can be expected to be important in both the sine and the cosine model. These expectations could be confirmed for the markers *PER2* and *PER3*: These genes had previously been reported to exhibit peak expression in the morning hours [36]. Indeed, for both markers we observed high, positive coefficients for the sine model and a smaller, negative (*PER3*) coefficient or a coefficient of 0 (*PER2*) for the cosine model. Similarly, the marker with the largest positive coefficient for the cosine model (*ENHO*, coefficient in the sine model: 0) had shown the highest expression values at 11 PM in our whole transcriptome sequencing dataset [16] and the marker with the largest negative coefficient for the cosine model (*UPB1*, coefficient in the sine model: 4.083) had previously shown the highest expression values at 8 AM in the morning [16].

Considering the top ten most important markers in both the sine and the cosine model, all markers except *ENHO* have previously been observed to exhibit diurnally rhythmic expression in human blood (cf. [16], Suppl. Table 8). *PER2* and *PER3* are well characterised core clock genes [37] (with *PER3* being the only high ranking marker in our model that had also been included in the model built by Lech et al. [12]). To the best of our knowledge, no direct function in the human body’s inner clock has been described for any of the other high-ranking markers, yet. However, several markers play a role in biological functions or pathways previously shown to be regulated by the body’s clock (e.g. innate and adaptive immune response [38] or metabolic functions [39]

A better understanding of the biological function of the markers identified as important in our time-of-day prediction model will be of general interest; yet it is not a pre-requisite for the inclusion of these markers in a model used in forensic applications.

### 3.5 Current limitations and future perspectives

The individuals participating in this study were chosen to represent an “average” of a healthy population. A detailed analysis of prediction performance for individual study participants showed that accuracies differed between individuals and were generally improved when accounting for interindividual differences in gene expression levels. Therefore, it can be concluded that knowledge of individual characteristics impacting prediction accuracies may help to build improved models in the future.

The regulation of the human body’s inner clock is highly complex. At the cellular level, the inner clock is constituted by a network of core clock genes which regulate the expression of downstream genes (“clock-controlled genes”) involved in different biological processes [40, 41]. Both the core clock genes as well as downstream pathways may be impacted by endogenous individual characteristics as well as exogenous factors, further increasing the complexity [41, 42]. A similarly complex biological phenomenon that has been extensively studied for application in forensic casework is the process of human aging. First studies to develop methods for predicting the age of the individual having deposited a forensically relevant biological trace focussed on the identification of markers (mostly differentially methylated DNA markers) and the establishment of prediction models for defined populations of healthy individuals [31, 35, 43–45]. These models have already been put to use as investigative tools in forensic casework. In several subsequent studies, researchers are now aiming to arrive at a better understanding of how individual endogenous characteristics such as sex and ethnicity as well as exogenous factors such as lifestyle and acquired diseases impact prediction accuracies and how these can be accounted for to improve general model performance [31, 46]. Thus, similar to the complex phenomenon of aging, the entirety of endogenous and exogenous factors impacting trace deposition timing exceeds the scope of a single study and further research to assess the impact of and possibility to account for these factors is required.

Apart from sample donors’ characteristics, the properties of the samples itself need to be taken into consideration when assessing the robustness of a prediction model. Blood traces analysed in forensic settings are commonly small in volume, dried and their integrity can be compromised by any number of detrimental environmental effects (such as exposure to light, moisture, microbial growth etc.).

While the transcriptome of a dried blood trace generally correlates with the transcriptome of liquid blood samples [47], the relative amount of individual markers can be altered by the process of drying and storage [11, 16]. Hence, the time-of-day prediction model in this study was built on gene expression measurements from blood traces that had been left to dry completely and all stored at room temperature for an identical time period (48 hours). The measurement results and prediction accuracies from these dried but fresh stains correlated well with results from stains stored for a longer time period (16 days). This is in accordance with previous studies showing that the transcriptome in whole blood remains relatively stable once the blood has completely dried [10, 11]. However, only a limited storage period and realistic but relatively favourable storage conditions (room temperature, dry, exposure to daylight) were tested experimentally in this study. As previous studies suggest that moist storage conditions allowing for microbial or fungal growth may drastically reduce nucleic acid integrity [11, 48], the impact of sample integrity/storage conditions on prediction accuracy needs to be assessed in more detail in future studies.

Additionally, forensically relevant blood traces may be of limited sample volume. In this study, all analyses have been performed with an RNA input amount of 25 ng. This amount of RNA was recoverable from dried blood in a volume of 50 µl, which is regularly encountered in casework scenarios. For lower input amounts, we observed a decrease in prediction accuracy. While a certain amount of total RNA will have to be available for analysis to be able to estimate the relative amount of individual transcripts present, it also needs to be taken into consideration that the targeted sequencing protocol using UMIs includes three bead purification steps (cf. Suppl. File 3), each likely leading to a loss of a certain sample proportion. Consequently, improved library preparation protocols may help to reduce sample loss thereby increasing method sensitivity.

Lastly, it needs to be highlighted that transcripts with rhythmic expression differ between cell types and biological fluids/tissues [49]. Thus, predictions obtained from the model presented in this study may only be considered reliable when the biological trace analysed contains whole blood from a single individual without cellular contributions from any other body fluid (which may be assessed by forensic STR typing and analysis of the expression of BFI markers included in the marker panel (cf. section 2.4).

For the purpose of this study, we have focussed on the analysis of mRNA markers, as this is currently the best explored type of RNA, it can be co-extracted with DNA from forensic traces using established protocols [23, 50] and its analysis requires only slight modifications to analysis procedures typically used for DNA thus allowing for fairly straightforward implementation into forensic molecular biology laboratory workflows. In future studies, other types of RNA such as miRNA and circRNA (which are relevant candidates in forensic studies due to their increased resistance to degradation [51–53]) might be explored. Besides RNA, other biomolecules such as metabolites [54–56] and hormones [57, 58] have been described to convey time-of-day information. Furthermore, here we focussed on time-of-day of trace deposition as only one aspect of forensic trace deposition timing. Complementing this information with insights on *time since deposition*, which is currently investigated by others [11, 59, 60], might lead to a more complete picture of this highly relevant contextualising aspect.

## 4. Conclusion

Our study provides the first prediction model for time-of-day of bloodstain deposition based on targeted RNA sequencing, demonstrating that time-of-day information is encoded in and may be recovered from blood traces as recovered from crime scenes. While the prediction accuracies of the method in its current state limit its use in the evaluative stage of a criminal trial, the method may nonetheless provide valuable information in the investigative phase. A better understanding of confounding endogenous as well as exogenous factors may lead to improved method performances in the future. In addition, the data presented in this study lay the groundwork for a better understanding and more standardised use of targeted cDNA sequencing for the purpose of gene expression quantification of medium and large sized marker panels in forensic applications.

Taken together, our study provides a relevant contribution to the growing knowledge on forensic trace contextualisation and represents an important step towards forensic trace deposition timing.

## 5. Material and Methods

### 5.1 Sample collection and preparation

#### 5.1.1 Study participants

Blood samples were obtained from 51 healthy volunteers (20 males, 31 females, median age: 26 years, range: 19-61 years). Volunteers with chronic or acute conditions known to affect regular sleep-wake cycles were excluded from the study (exclusion criteria: irregular sleep-wake-rhythms (e.g. jet-lag, shiftwork), diagnosed sleeping disorders, pregnancy or lactation, diseases of the hematopoietic system (current or within the last five years), use of medication known to interact with the diurnal rhythm (such as hormone preparations) and blood thinners (current or within the last year)). The study participants’ chronotypes were assessed using the German version of the Horne-Ostberg-Morningness-Eveningness-Questionnaire D-MEQ [61]. Study participant characteristics are summarised in Suppl. Table 4.

#### 5.1.2 Collection of blood samples

Collection of blood samples was performed at eight specified time points (8 AM, 11 AM, 2 PM, 5 PM, 8 PM, 11 PM, 2AM, 5 AM) over the course of an entire day. Donors were instructed to refrain from the intake of alcohol and other drugs on the entire day of sample collection and from sleeping between the first (8 AM) and last time point of sample collection (5 AM) but otherwise pursued regular every-day activities. For individual samples (n = 10), donors indicated having deposited samples up to 30 minutes before or after the specified time point (Suppl. Table 3); however, for the purpose of this study these were treated as having been deposited at the exact specified timepoint. Sample donors filled in a questionnaire on time of food consumption, intake of medication and performing intense physical activities on the day of sample collection.

Blood samples were self-collected by the volunteers via fingerprick with sterile lancets (28 G SafetyLancets (MyLife, Ningbo Medsun Medical Co., Zhenhai, Ningbo, China, Ref. 7100031) or 21 G Safety Lancets (Vivomed GmBH, Geislingen, Germany, Ref. 19170010)). 50 µl fingerprick blood was collected with the help of untreated “neutral” Minivettes (Sarstedt, Nürmbrecht, Germany, Cat. No. 17.2111.050). (A detailed description of the sample collection procedures is provided in Supplementary Figure 1 of [16]).

Each donor collected two samples of 50 µl per time point. The blood was immediately transferred to clean 2.0 ml plastic tubes (Safe-Lock Tubes, Forensic DNA Grade (Eppendorf, Hamburg, Germany, Catalogue number: 0030123620)) in which the samples were left to dry at room temperature by leaving tube caps open. Samples were picked up and transported to the research facility on the day following sample collection. For the period of transportation, tube caps were briefly closed (for a maximum period of 2.5 hours) and then further let dry with open caps at the research facility. Sample tubes were closed and samples transferred to -80°C for prolonged storage after a drying time of exactly 48 hours or 16 days (for one of the two samples collected per participant respectively). A detailed overview of all samples collected and the individual sample characteristics is provided in Suppl. Table 3.

Regarding individual characteristics and activities, it remains to be pointed out that all data presented in this study was self-reported by study participants. While all study participants (or at least one individual for groups of study participants) were encountered by a researcher in person and were generally considered as trustworthy, it cannot fully be ruled out that individual statements (e.g. regarding medication or adherence to the study protocol) were incomplete or erroneous.

#### 5.1.3 RNA isolation and quantification

For all samples, total RNA was extracted using the mirVana miRNA Isolation Kit (Thermo Fisher Scientific, Massachusetts, USA, Cat. No. AM1560/ AM1561). For sample lysis, samples were incubated in 450 µl Lysis/Binding Buffer complemented with 30 µl Proteinase K (Roche Diagnostics, Mannheim, Germany, Ref.: 03115828001) at 56°C for 30 minutes. RNA was subsequently isolated from the lysate following manufacturer’s instructions (“Total RNAs” protocol) and eluted in a final volume of 35 µl nuclease-free water (Invitrogen, Carlsbad, USA, Ref.: 10977035), which was added to the filter twice. Residual genomic DNA was removed using the TURBO DNAse free kit (Thermo Fisher Scientific, Cat. No. AM1907) with 2 µl DNase per sample as per manufacturer’s instructions. RNA concentrations were determined fluorometrically using the Qubit RNA HS Kit (Thermo Fisher Scientific, Cat. No. Q32855) on a Qubit 2.0 device (Thermo Fisher Scientific).

### 5.2 Targeted gene expression quantification

Expression quantification of target genes was to be performed by targeted cDNA sequencing on an Ion S5 benchtop instrument. As the inclusion of Unique Molecular Identifiers (UMIs) had proven to significantly increase gene expression quantification accuracies in a small-scale experiment performed prior to this study [25], the QIAseq targeted RNA Panels kit (QIAGEN, Hilden, Germany, Cat. No. 333005) was chosen to be used in this study.

#### 5.2.1 Marker selection and panel design

The targeted sequencing panel was constructed to include four types of markers: time-of-day candidate predictors, reference gene candidates, markers for body fluid identification and gDNA control markers. **Time-of-day candidate predictors** had previously been identified by whole transcriptome sequencing in [16]. Of the 81 candidate predictors identified in this study, five were small RNAs (*MIR6883, MIR223, MIR7848, RNY1, SNORA73A*). As these targets would have required a different approach for targeted sequencing, they were not included in this study. In addition to the remaining 76 candidate predictors identified by whole transcriptome sequencing, eight candidate predictors (*AVIL, DAAM2, HNRNPDL, MKNK2, NR1D1, NR1D2, PER2, PER3*) were included in this study that had not met all selection criteria in the previous whole transcriptome sequencing study, but have been described as rhythmically expressed in a relevant number of previously published studies/datasets (appearing in ≥ seven previous publications/datasets as summarised in Supplementary Table 8 in [16]). Nine **reference gene candidates** were included based on selection criteria described in [16], section 2.2.6. To be able to identify **body fluids** present in the sample, previously selected and validated mRNA markers for the identification of blood, saliva, semen, skin, vaginal secretions and menstrual blood [18] were included in the panel. Following a manufacturer’s recommendation, six **gDNA controls** allowing to assess the presence of human genomic DNA (gDNA) in the samples were additionally included in the panel.

Taken together, the initial version of the primer panel contained primers for a total of 122 markers. After initial performance testing on 44 samples (data from 11 individuals taken at time points 8 AM, 2 PM, 8 PM and 2 AM), a total of 10 candidate predictors were excluded due to low expression (*ST20, TSHZ2*), high expression (*DUSP1*) or lack of statistically significant expression differences (*CEBPD, DDIT4, IL23A, KCNE3, RELL1, TNFSF14, TXN*), (data not shown). All further samples were thus processed using a panel of 112 markers, including six gDNA control markers, nine reference gene candidates, 23 genes for the identification of body fluids present and 74 candidate genes for time-of-day prediction (Suppl. Table 5).

Primers for all selected markers were custom designed in cooperation with the QIAGEN Enterprise Genomic Solutions team to specifically amplify selected targets and minimise potential for primer-primer interactions. A cDNA-specific design was possible for all but two targets (*LCE1C* and *GCNT4*) by choosing primer binding sites on two different exons. Interspersed intronic sequences were chosen to be of sufficient length for the corresponding amplicon resulting from genomic DNA to be produced less efficiently and additionally to be removed from the sample in the bead-based size selection step. The primer sequences are proprietary, but primers can be reordered from QIAGEN by reference to the panel design number CRHS-18209Z-112.

#### 5.2.2 Library preparation using the QIAseq Targeted RNA panel

Targeted sequencing libraries were prepared from 25 ng total RNA per sample using the QIAseq Targeted RNA Panel (Cat. No. 333005, QIAGEN) according to the manufacturer’s “QIAseq Targeted RNA panels for Ultra-Low Input and FFPE Samples v2” protocol, which is summarised in Suppl. File 3. For the dilution study, libraries from 12 samples from three randomly selected individuals (v030, v031, v071) at four timepoints (8 AM, 2 PM, 8 PM, 2 AM) were additionally prepared from 5 and 0.5 ng total RNA input.

For sequencing on an Ion GeneStudio S5 benchtop sequencer, libraries were indexed with the QIAseq Targeted RNA 96-Index HT L for Ion Torrent Kit (Cat. No. 333217, QIAGEN) in the Second Stage Universal Index PCR step. Final sequencing libraries were quantified via qPCR using the QIAseq Library Quant Assay Kit configured for Ion Torrent (Cat. No. 333314, QIAGEN) with ½ reaction volumes on a 7500 Fast Real-time PCR (Thermo Fisher Scientific) or a QuantStudio 5 instrument (Thermo Fisher Scientific).

A total of 532 sequencing libraries were prepared in this study. Excluding libraries prepared from diluted samples, final libraries had an average concentration of 39,970 pM with a relative standard deviation of 0.36. For only four out of 532 samples, library preparation failed initially and libraries needed to be prepared again. Hence, the overall library preparation process was considered robust and consistent.

#### 5.2.3 Targeted Sequencing on an Ion S5 benchtop sequencer

Targeted sequencing was performed on an Ion GeneStudio S5 benchtop sequencer (Thermo Fisher Scientific). For this purpose, sequencing libraries were normalised to a concentration of 50 pM in nuclease-free water (libraries with concentrations < 50 pM were prepared again). Sequencing libraries with differing index primers were pooled for combined sequencing of 29-30 samples on one Ion 530 sequencing chip (Cat. No. A27764, Thermo Fisher Scientific). For UMI-based sequencing libraries, sufficient sequencing depth is necessary to accurately quantify the number of different UMIs per target marker present in each sample. Details regarding sequencing depth optimisation are provided in Suppl. File 2. Templating and sequencing were performed with the Ion S5™ Precision ID Chef & Sequencing Kit (Thermo Fisher Scientific, Cat. No. A33208) on an Ion Chef and an Ion GeneStudio S5 benchtop sequencer (both Thermo Fisher Scientific) respectively. Templating was performed with default settings and the “HID Ion S5 Chef Templating 02” protocol. Default sequencing settings were used with 500 flows per sequencing chip.

#### 5.2.4 Bioinformatic processing of sequencing data

Raw data processing and demultiplexing was performed in the Ion Torrent Suite (v. 5.10.1) using default analysis settings without a reference library and IonXpress barcode settings. Demultiplexed raw sequencing files (.basecaller.bam-files) were exported and further processed using the “QIAseq Targeted RNA panels” workflow with “custom catalog #” targets on the GeneGlobe platform (QIAGEN). Briefly, during this workflow, sequencing reads are trimmed, quality filtered and separated into target and UMI sequence. Target sequences are then aligned to the GRCh38 reference genome using the STAR RNA read mapper [62]. Target read counts are converted to UMI read counts by clustering target reads with identical UMI sequences [63]. Nearly identical UMI counts were obtained when processing raw data files with the “Quantify QIAseq RNA Workflow” on the CLC Genomics Workbench (Version 23.0.4, QIAGEN Aarhus A/S), (data not shown), thus the more streamlined GeneGlobe workflow was deemed suitable for the purpose of this project.

To obtain reliable UMI quantification results, we aimed for a minimum number of reads used for UMI counting of ∼400.000 per sample (cf. Suppl. File 2). No relevant gDNA contamination was observed (counts in gDNA controls ≤1).

### 5.3 Statistical Analysis

All statistical analyses were performed with the statistics software R [64]. All tests were two-side with a significance level of 0.05.

#### 5.3.1 Read count normalisation

Gene expression quantification methods require normalisation procedures to account for technical variation in the various sample preparation steps. While robust normalisation procedures have been established and guidelines have been proposed for the analysis of qPCR data [65, 66], dPCR data [67], microarray experiments [68] as well as whole transcriptome sequencing approaches [69], to the best of our knowledge no such standards or guidelines have been established for the analysis of data from targeted sequencing of cDNA.

We thus decided to apply and compare six different normalisation procedures (which have been applied in previous studies employing targeted RNA sequencing or adapted from other technical approaches such as qPCR or whole transcriptome sequencing). A detailed description of these comparisons is provided in Suppl. File 1. Briefly summarised, all tested normalisation procedures were suitable to decrease variability between groups (i.e. samples collected at different timepoints) as compared to non-normalised data, while maintaining or increasing variability between groups and may thus be considered suitable normalisation procedures. While smaller differences were observed, all six normalisation procedures showed the same general trends. We therefore concluded that while the inclusion of a normalisation step is essential for accurate gene expression quantification, different normalisation approaches may lead to comparable results. Consequently, all downstream analyses were performed using two different normalisation procedures (chosen based on generalised η^2^-squared values, cf. Suppl. File 1).

*CPM (Counts Per Million) normalisation:* For each sample, the raw read count of each target gene was divided by the total read count of all targeted genes and multiplied by one million. CPM thus represent the number of counts expected to be observed for the respective target gene if each library was sequenced to a total of exactly one million reads.

*"Refall" normalisation*: For each sample, the raw read count of each target gene was divided by the average read counts of all reference gene candidates in that same sample. (Normalisation to only a subset of reference gene candidates gave similar, but not better results (cf. Suppl. File 1)).

Raw and normalised read counts are provided in Suppl. Table 6.

#### 5.3.2 Initial evaluation of marker suitability

Based on the results of the rarefaction analysis (Suppl. File 2), expression quantification of the markers *ATG2A* and *IRS2* was not considered reliable and the markers were excluded from any further analysis. For all remaining 72 time-of-day candidate markers, we checked for timepoint dependence of individual marker expression. First, we removed those markers which showed no significant difference between the eight time points (Friedman Test, p-value > 0.05, Suppl. Table 7). Furthermore, we calculated a mixed linear model with expression as outcome, time point as fixed effect and individual as random effect and compared that to a linear model without the random effect and to a mixed linear model without the fixed effect (using the *lmer()* function from the R package „*lme4*“ [70]). The models were compared with the *anova* function and markers were excluded if either of these comparisons was not significant (Suppl. Table 7). This approach led to the removal of two markers from the CPM-normalised dataset *(IL18R1, GABARAPL1*), whereas no marker was removed from the Refall-normalised dataset. Afterwards, we calculated the Pearson correlations between all remaining time-of-day candidate prediction markers for each timepoint (Suppl. Fig. 5). For all marker pairs with Pearson correlation above 0.80 for all timepoints, we removed the marker with the higher p-value of the aforementioned Friedman test, leading to the removal of three markers from the Refall-normalised dataset (*REPS2, MEGF9, TBXAS1*) and one marker from the CPM-normalised dataset (*TRABD2A*). After these initial evaluation steps, a total of 69 markers remained in both the Refall- and the CPM-normalised dataset.

#### 5.3.3 Modelling time-of-day

Due to the cyclic nature of time-of-day, commonly used and well-established regression models are not directly applicable to this variable. We thus employed two different approaches for modelling time-of-day:

a. Following an approach described previously [17], we obtained variables that can be handled with common regression models by considering time as the angle of the hour hand of a 24-hour clock and subsequently trigonometrically transforming this angle to a sine and a cosine component (Figure 1). Separate regression models were then constructed for each component and the predictions obtained from the separate sine and cosine models were re-transformed to obtain an angle (corresponding to a specific time point on a 24-hour clock).
b. Alternatively, time-of-day was modelled categorically with each sample deposition time considered as one category (resulting in a total of eight categories).

#### 5.3.4 Assessing model performance

To predict time-of-day based on gene expression, different statistical approaches were applied, individually optimised as described below and compared with regards to root mean squared error (RMSE). Additionally, we calculated the accuracy per category (within +/- 1.5 hours), accuracy within +/- 4 hours (this range was chosen as it was considered “useful” for a typical forensic casework constellation, albeit being somewhat arbitrary) and the mean absolute error. A direct comparison of the continuous and the categorical modelling approaches of time of day is not straightforward as different performance metrics are better suitable to evaluate either categorical or continuous predictions. Due to the continuous nature of time-of-day, RMSE was considered as the most relevant performance metrics for comparative purposes in this case.

Model performance was assessed using five-fold cross-validation. For this purpose, the dataset was split into five groups. To ensure that training and test sets were truly independent, samples from one individual were kept in a single group, so that models would never be tested on samples from an individual from who another sample had been part of the training set. Groups were set up with a balanced distribution of participants’ age and sex. Samples in each group were subsequently split into eight subgroups so that each sample from a single individual was part of a different sub-group and the time-points were evenly distributed across sub-groups. Cross-validation was then performed by training a model on four of the five groups and obtaining predictions for the remaining group as test set. This was repeated four times so that each sample was part of the test set once and all test set predictions were combined to calculate model performance metrics. To ensure comparability between models, identical groups and sub-groups (shown in Suppl. Table 3) were used for each of the prediction algorithms.

#### 5.3.5 Prediction algorithms

Four different prediction algorithms were tested and compared: Random Forest (RF), Support Vector Machines (SVM), Penalised Regression (PR) and the previously published algorithm TimeSignatR (which is also based on penalised regression). These approaches are typically suitable for situations like the one at hand, where many influence variables are considered and the number of samples is relatively small. Other conceivable algorithms, such as multinomial regression, linear regression without penalisation, gradient boosting and neural networks, proved to perform poorly during preliminary testing (data not shown). This was to be expected, since without penalisation too many variables remain in the regression and neural networks usually require large sample sizes. Furthermore, we decided against the use of mixed models, as only a single sample per individual would usually be available for our use case.

RF and SVM were applied for both the continuous and the categorical parametrisations of time-of-day whereas PR and TimeSignatR were only applied to the continuous parametrisation. For the discrete version of the PR approach, the eight classes for the categorical outcome led to convergence issues and the TimeSignatR algorithms was explicitly designed only for continuous outcomes. Due to the sine and cosine representation of time of day in the continuous approach, two models, each for every component, have to be estimated independently. In the categorical parametrisation, in contrast, only a single model is estimated. Each algorithm was optimised individually as described below.

##### Random Forest (RF)

We used the R package *ranger* [71] for random forest modelling for the continuous and the categorical outcomes. Based on pretesting, the number of trees was set to 100. We ran separate cross-validations for different number of variables to possibly split at each node (parameter *“*mtry*“*) (for multiples of five). The best *“*mtry*“*-value was chosen based on the best RMSE for all time points of the respective cross-validation result.

##### Support Vector Machine (SVM)

SVMs were implemented by using the R package *e1071* [72]. We ran separate cross-validations for each choice of kernel function (linear, polynomial, radial, sigmoid). The best kernel function was chosen based on the best RMSE for all time points of the respective cross-validation result.

During each cross-validation step, the respective hyperparameters for the kernel functions were optimised by grid tuning, an inner cross-validation. Here, each of the remaining four groups from the training dataset served in turn as test set for the models trained by the other three groups across a grid of all relevant hyperparameters. Hyperparameters being used were cost (all kernel functions), epsilon (all kernel functions), gamma (polynomial, radial and sigmoid kernel), coef0 (polynomial and sigmoid kernel) and degree (polynomial kernel). The set of hyperparameters were chosen which yielded the model with the lowest error according to e1071’s *tune()*-function (classification error for categorical time-of-day, mean squared error for continuous time-of-day) across the inner cross-validation on the four training groups.

##### Penalised Regression (PR)

The R package *glmnet* [73] was used to implement the penalised regression approach. During the five-fold cross-validation, the hyperparameters alpha and lambda of each model were optimised by grid tuning. Within the respective training set, an additional five-fold cross-validation was applied on a grid of alpha and lambda values and the parameter set which minimised the mean cross-validated error (default measure of the *cv.glmnet()-*function) was chosen.

##### TimeSignatR

TimeSignatR was performed as described in [17] using adapted versions of the R scripts available from github.com/braunr/TimeSignatR (last access: 2025-01-31). Briefly summarised, a multi-response penalised regression model (*family: “mgaussian”*) was trained for the sine and cosine components of time-of-day as response variables using the *glmnet* R package [73] (v. 4.1-8). The model was trained on Refall-/CPM-normalised data omitting the within-subject renormalisation step included in [17]. For each model trained in five-fold cross-validation, hyperparameters alpha and lambda were individually optimised using grid-tuning and ten-fold cross-validation (within the respective training set).

#### 5.3.6 Relationship between prediction errors (PR) and per category prediction probabilities (SVM)

To assess the informative value of per category prediction probabilities (obtained from the categorical SVM model), the relationship between prediction errors (absolute difference between true and predicted time of deposition) from the PR model and the prediction probabilities obtained for the corresponding category was evaluated. As QQplots generated with the *ggpubr* R package [74] suggested non-normal distribution of both variables, a Spearman’s rank correlation test was calculated. Test results are indicated with the subscript “Spearman”.

#### 5.3.6 Within-subject renormalisation

Within-subject renormalisation was performed as described in [17]: For each gene, the mean Refall-normalised expression was calculated per individual, and the individual Refall-normalised expression values were then normalised to the per-subject mean, so that the within-subject renormalised values indicate the deviation from the individual expression mean over time. Model training and testing was then performed as described in section 5.3.4 and 5.3.5.

#### 5.3.7 Extending the prediction model to include individual characteristics

It is to be expected that other variables also correlate with the diurnal rhythm. We therefore examined how the predictive quality of the models changes when age and sex of the subjects are included in the modelling in addition to gene expressions. Here, age was given as the age at sampling (in years) and sex as dichotomous variable. We used the same procedure as above to cross-validate the algorithms and generate the respective final PR models (section 5.3.4 and 5.3.5).

#### 5.3.8 Assessing the impact of individual characteristics on prediction error

The impact of individual characteristics and activities on model prediction error was evaluated by mixed-effects models with prediction error as outcome and random intercepts for the individuals using the *lme4* [70] and *lmerTest* [75] packages in R. Age and sex were combined into a single model, but due to the exploratory nature of all further analyses, additional characteristics and activities were tested in separate models. Model assumptions including normal distribution of residuals and homogeneity of variance between the levels of a predictor were assessed using functions provided in the *DHARMa* R package [76] and met for all models tested after log(x+1) transformation of the prediction error. Test results are indicated with the subscript “LMM”. Due to the absence of statistically significant results, multiple testing corrections were not performed.

### 5.4 Assessing the impact of sample conditions

#### 5.4.1 Storage duration

Each study participants had deposited two blood samples per deposition timepoint. These were then stored at room temperature for exactly 48 hours and 16 days respectively. To estimate the impact of sample storage duration, samples from 25 randomly selected individuals (v001, v002, v009, v010, v011, v013, v015, v016, v017, v018, v021, v022, v031, v035, v039, v040, v045 v059, v063, v067, v069, v070, v071, v072 and v073) collected at four different timepoints (8 AM, 2 PM, 8 PM and 2 AM) stored for 16 days at room temperature were extracted and analysed as described in sections 5.1.3 and 5.2.

For direct comparison, a PR model as described in section 5.3.5 was trained on data from 208 samples (eight timepoints) from the remaining 26 individuals and subsequently tested on 200 samples (four timepoints) from the above-described individuals, both after storage for 48 h and 16 d (100 samples each). The resulting predictions for the sine and cosine components of time-of-day were directly compared. As QQplots generated with the *ggpubr* R package [74] suggested normal distribution of both variables, a Pearson correlation test was performed using the *cor.test* function from the *stats* R package [64].

#### 5.4.2 Sample dilution

To estimate the impact of reduced RNA input to library preparation, samples from three randomly selected individuals (v030, v031, v071), collected at four different timepoints (8 AM, 2 PM, 8 PM and 2 AM) were additionally subjected to library preparation with sub-optimal input amounts of 5 ng and 0.5 ng (20 and 2 % of t e optimal input amount respectively .

For direct comparison, a PR model as described in section 5.3.5 was trained on data from 384 samples (eight timepoints) from the remaining 48 individuals and subsequently tested on 12 samples (four timepoints) from the above-described individuals, with all three input amounts. The resulting predictions for the sine and cosine components of time-of-day for the 5 ng and 0.5 ng dilutions were directly compared to the undiluted samples. As QQplots generated with the *ggpubr* R package [74] suggested normal distribution of both variables, a Pearson correlation test was performed using the *cor.test* function from the *stats* R package [64].

## 6. Supplementary Materials

Suppl. File 1: Comparison of normalisation procedures for targeted RNA sequencing data

Suppl. File 2: Optimisation of sequencing depth

Suppl. File 3: Library preparation QIAseq targeted RNA sequencing with UMIs

Suppl. Figure 1: Pairwise correlation of the absolute prediction errors obtained for each of the four continuous prediction models

Suppl. Figure 2: Prediction error and predicted probability shown in relation to donor characteristics and activities

Suppl. Figure 3: Scatterplot showing true and predicted times for individuals according to match group

Suppl. Figure 4: Boxplot showing Refall-normalised expression of BFI markers in blood samples

Suppl. Figure 5: Correlation and selection of markers

Suppl. Table 1: Comparison of model performances for four modelling algorithms Suppl. Table 2: Individual prediction results for the final model

Suppl. Table 3: Information on individual samples Suppl. Table 4: Information on study participants

Suppl. Table 5: List of markers targeted with the final primer panel

Suppl. Table 6a: Raw read counts (for all samples sequenced in this study)

Suppl. Table 6b: Refall-normalised read counts (for all samples sequenced in this study) Suppl. Table 6c: CPM-normalised read counts (for all fresh (dried for 48 h), undiluted samples sequenced in this study)

Suppl. Table 7a: Timepoint dependencies for the Refall-normalised data

Suppl. Table 7b: Timepoint dependencies for the CPM-normalised data

## 7. Declarations

### 7.1 Ethics approval and consent to participate

All sample donors provided written informed consent. The study protocol was reviewed and approved by the ethics committee of the University Hospital Cologne.

### 7.2 Consent for publication

Not applicable.

### 7.3 Availability of data and materials

The raw sequencing data files generated for the current study are not publicly available due to the sensitive nature of the data but are available from the corresponding author for the sole purpose of replicating the procedures and results presented in the article and providing that the data transfer and storage is in agreement with European Union legislation on the general data protection regulation.

All other data generated or analysed during this study are included in this published article and its supplementary information files.

### 7.4 Competing interests

The authors declare that they have no competing interests.

### 7.5 Funding

The study was funded by the Deutsche Forschungsgemeinschaft (Project number 453376443).

The funders of the study had no role in study design, data collection, analysis and interpretation, or writing of the manuscript.

### 7.6 Authors’ contributions

AG and CC conceptualised the project and acquired funding; AG collected samples and performed laboratory experiments; AG and SS analysed data and visualised results; CC and AC supervised data analysis and interpretation; AG wrote the initial manuscript draft; all authors reviewed, edited and approved the final manuscript.

## 7.8 Acknowledgements

We sincerely thank all anonymous sample donors for taking part in this study. Additionally, we would like to thank Christian Klug and the company QIAGEN for their great support during the establishment of the targeted cDNA sequencing assay and Maximilian Neis (University Hospital Cologne) for his introduction to the Ion S5 benchtop sequencing instrument.

## Notes

### Competing Interest Statement

The authors have declared no competing interest.

