## Supplementary Figures for "TrACES of Time: Towards estimating time-of-day of bloodstain deposition by targeted RNA sequencing"

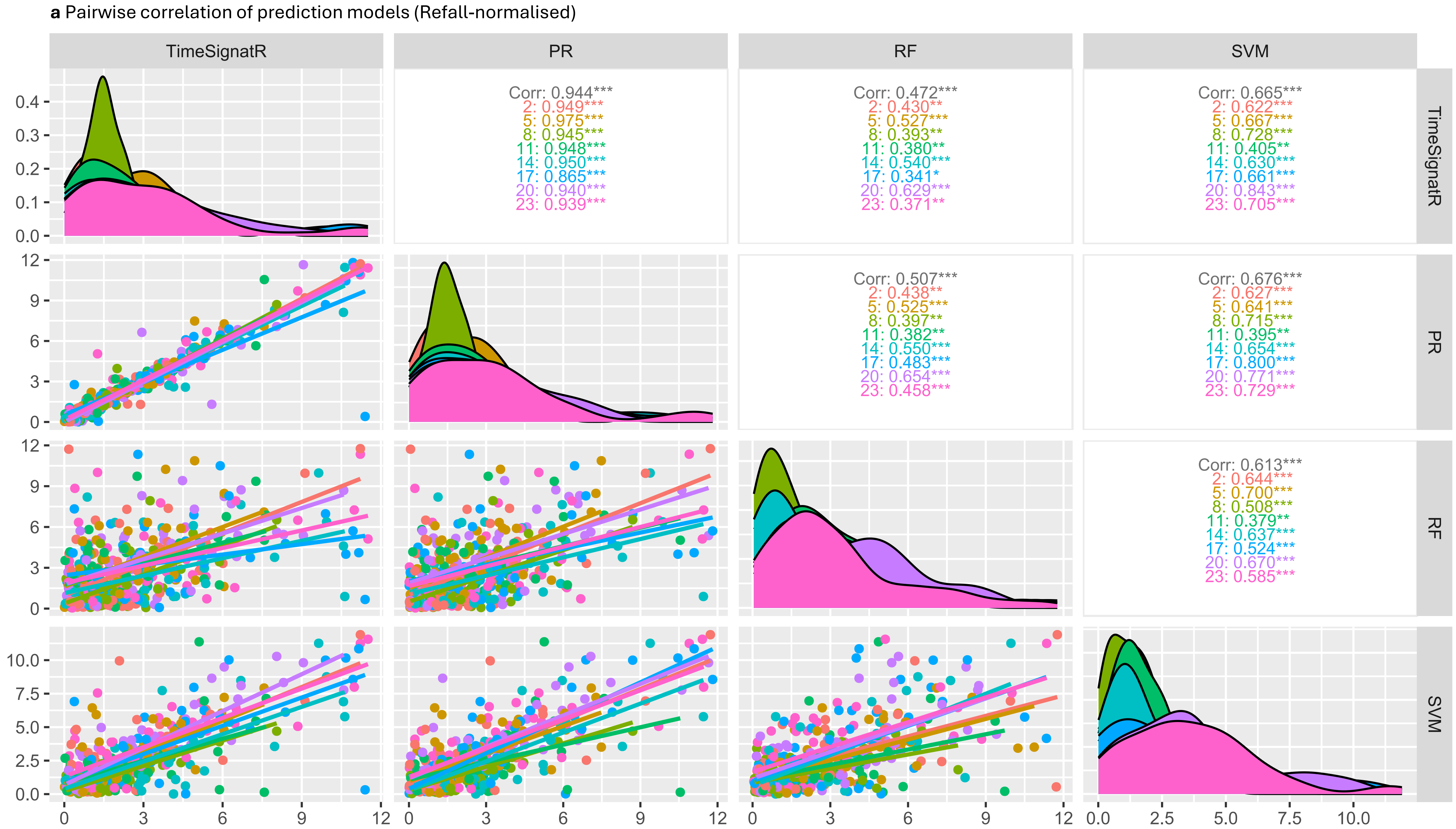


**Suppl. Figure 1a: Pairwise correlation of the absolute prediction errors obtained for each of the four continuous prediction models** obtained by five-fold cross-validation (n=408 samples from 51 individuals deposited at eight different timepoints, dried for 48 hours; Refall-normalised dataset).
Lower panels show the pairwise comparison of individual prediction errors (x-axis: True time in hours, y-axis: prediction error in hours); upper panels show the Spearman correlation coefficient (overall and per deposition timepoint) and diagonal panels show the distribution of absolute prediction errors (x-axis: prediction error in hours, y-axis: density).

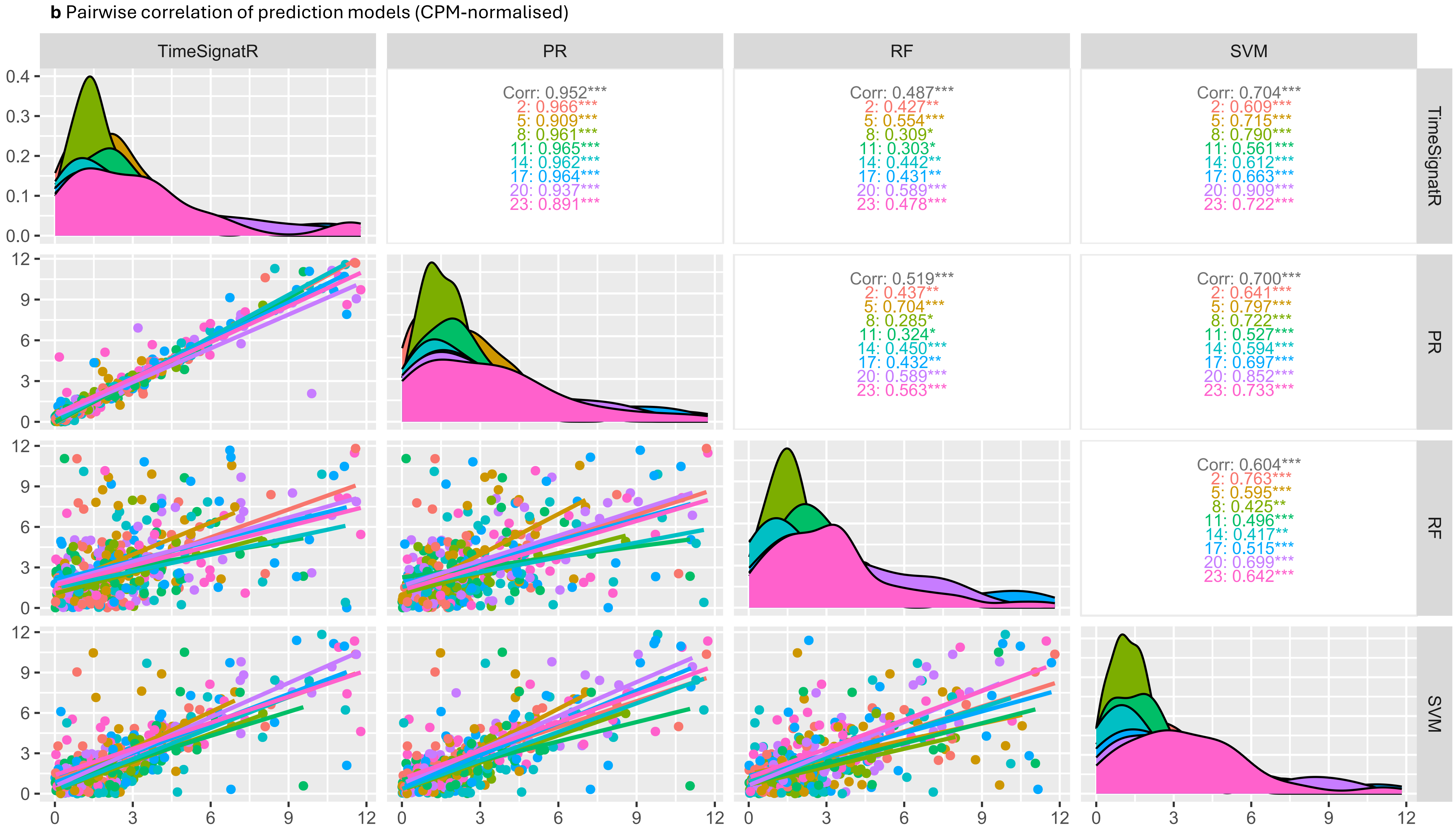


**Suppl. Figure 1b: Pairwise correlation of the absolute prediction errors obtained for each of the four continuous prediction models** obtained by five-fold cross-validation (n=408 samples from 51 individuals deposited at eight different timepoints, dried for 48 hours; CPM-normalised dataset).
Lower panels show the pairwise comparison of individual prediction errors (x-axis: True time in hours, y-axis: prediction error in hours); upper panels show the Spearman correlation coefficient (overall and per deposition timepoint) and diagonal panels show the distribution of absolute prediction errors (x-axis: prediction error in hours, y-axis: density).

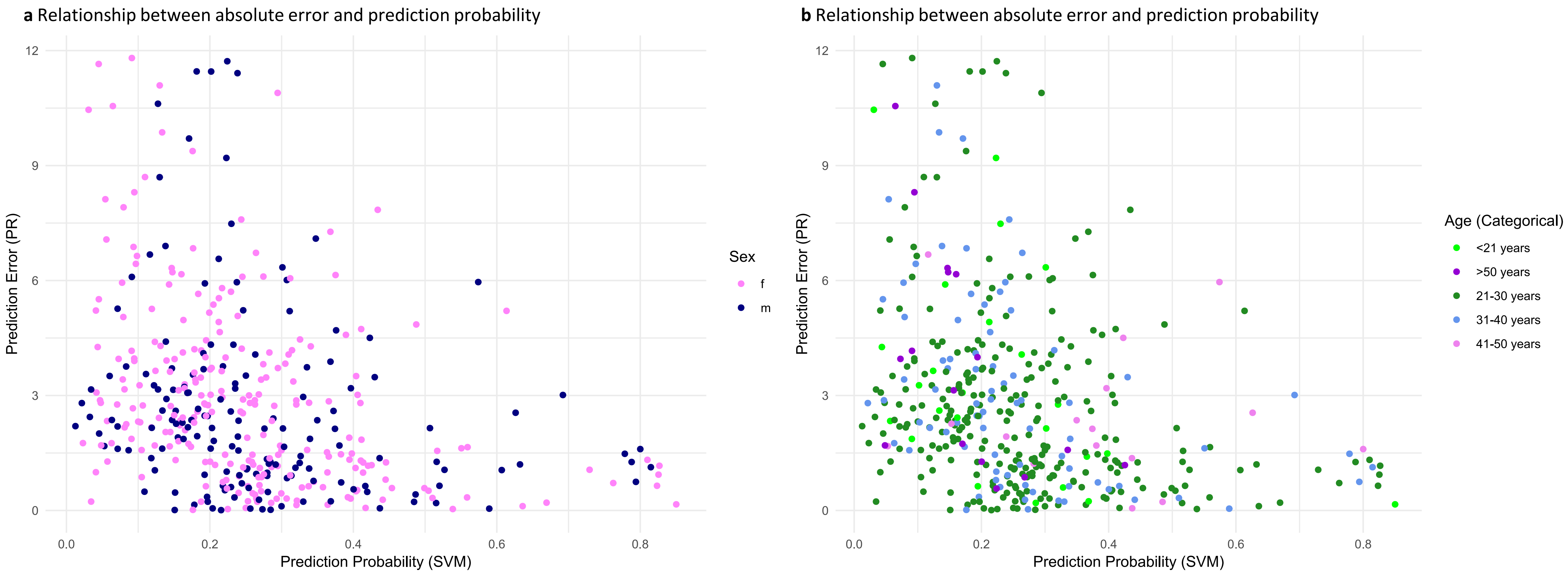


**Suppl. Figure 2: Prediction error and predicted probability shown in relation to donor characteristics and activities:**
Predictions obtained for samples in the respective test set of five-fold cross-validation (n=408 samples from 51 individuals deposited at eight different timepoints, dried for 48 hours; Refall normalised dataset)
**a:** Donor sex, **b:** Donor age (categorical) , **c:** Donor chronotype (assessed using the D-MEQ questionnaire), **d:** Time of the year of sample deposition, **e:** Consumption of caffeine (within 3 hours prior to sample deposition) , **f:** Deviations from study protocol instructions (consumption of alcohol or napping within 3 hours prior to sample deposition) , **g:** Individuals taking medication (medication indicated in brackets (vitamins and oral contraceptives not included)).

c Relationship between absolute error and prediction probability

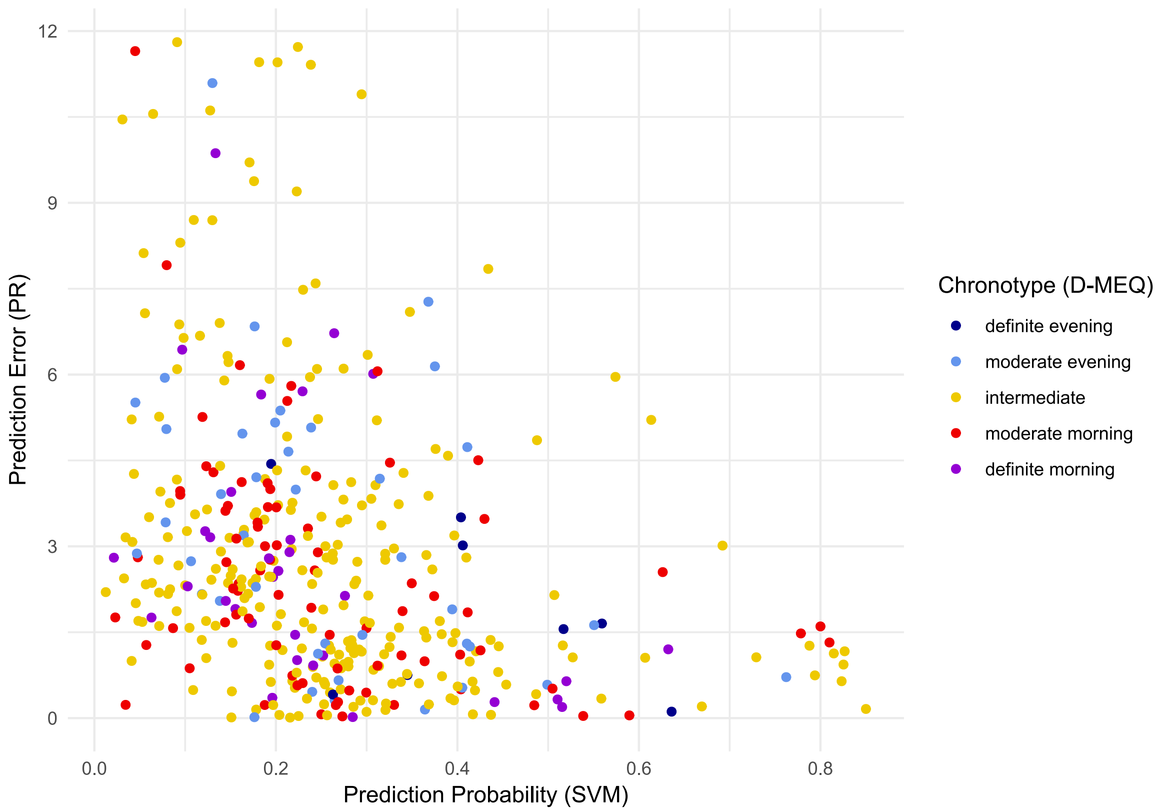


d Relationship between absolute error and prediction probability

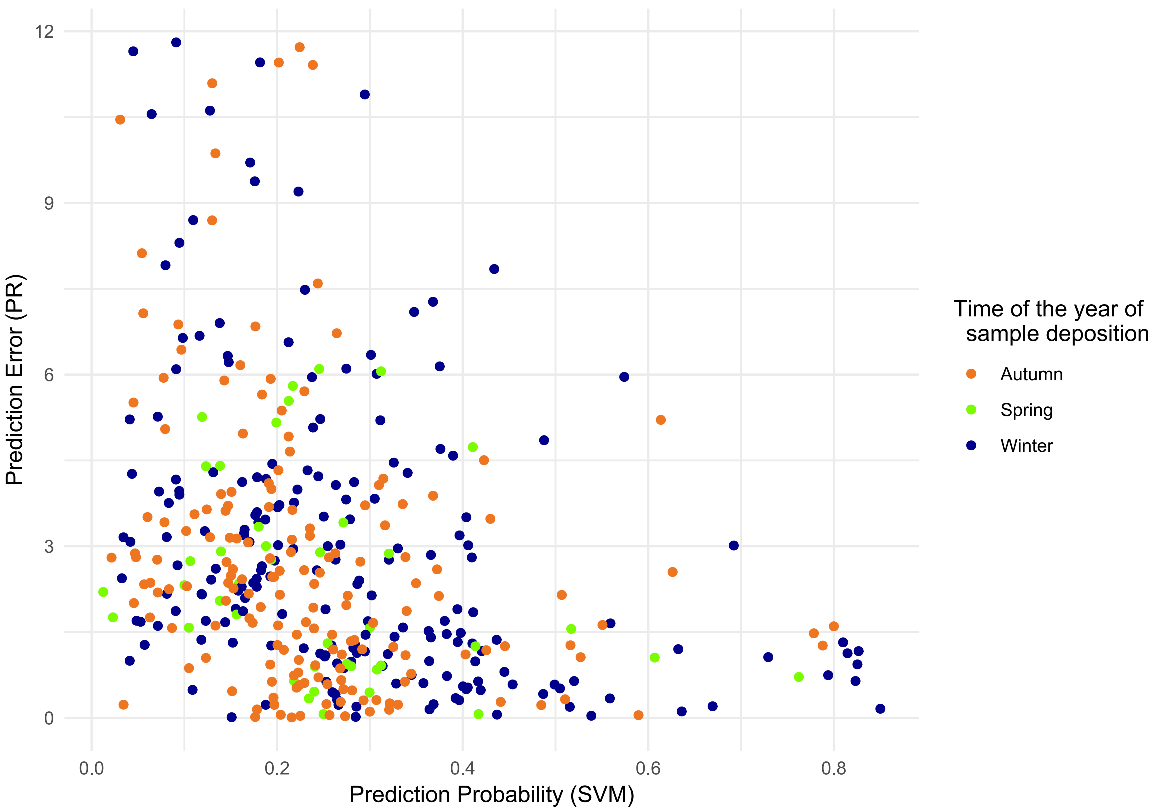


**Suppl. Figure 2: Prediction error and predicted probability shown in relation to donor characteristics and activities:**
Predictions obtained for samples in the respective test set of five-fold cross-validation (n=408 samples from 51 individuals deposited at eight different timepoints, dried for 48 hours; Refall normalised dataset)
**a:** Donor sex, **b:** Donor age (categorical) , **c:** Donor chronotype (assessed using the D-MEQ questionnaire), **d:** Time of the year of sample deposition, **e:** Consumption of caffeine (within 3 hours prior to sample deposition) , **f:** Deviations from study protocol instructions (consumption of alcohol or napping within 3 hours prior to sample deposition) , **g:** Individuals taking medication (medication indicated in brackets (vitamins and oral contraceptives not included)).

**e** Relationship between absolute error and prediction probability

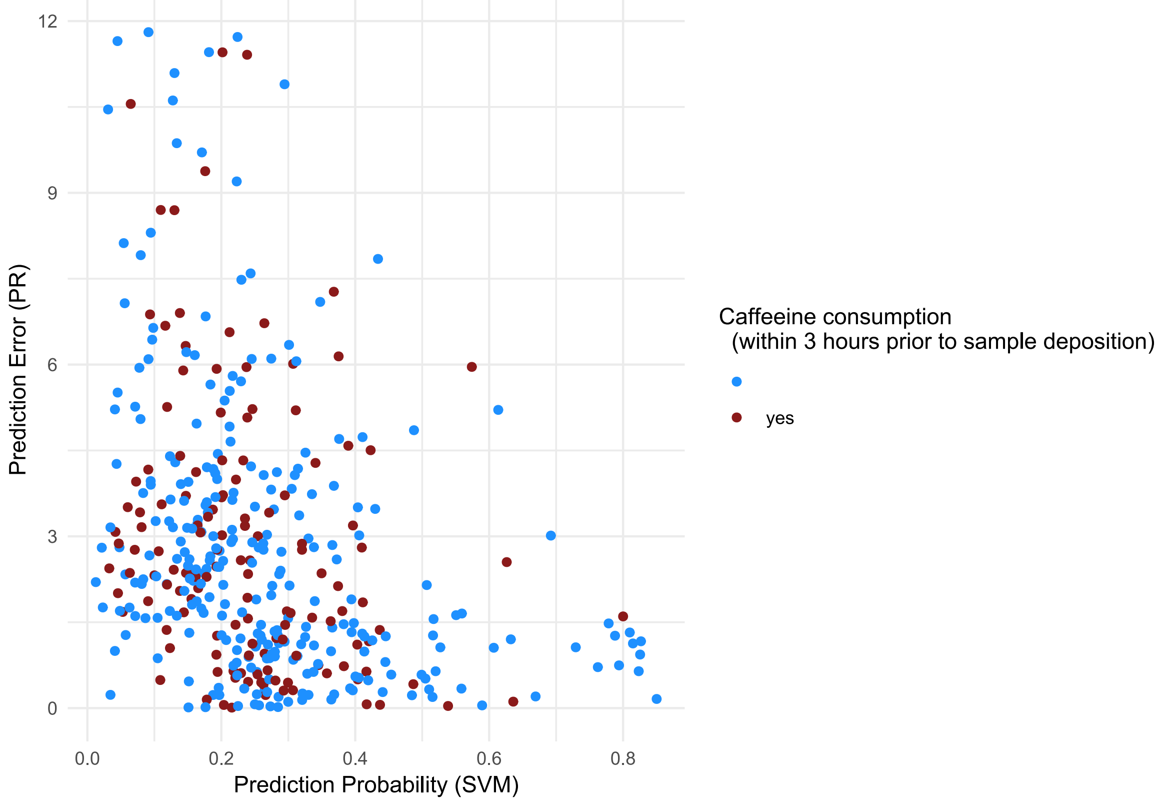


**f** Relationship between absolute error and prediction probability

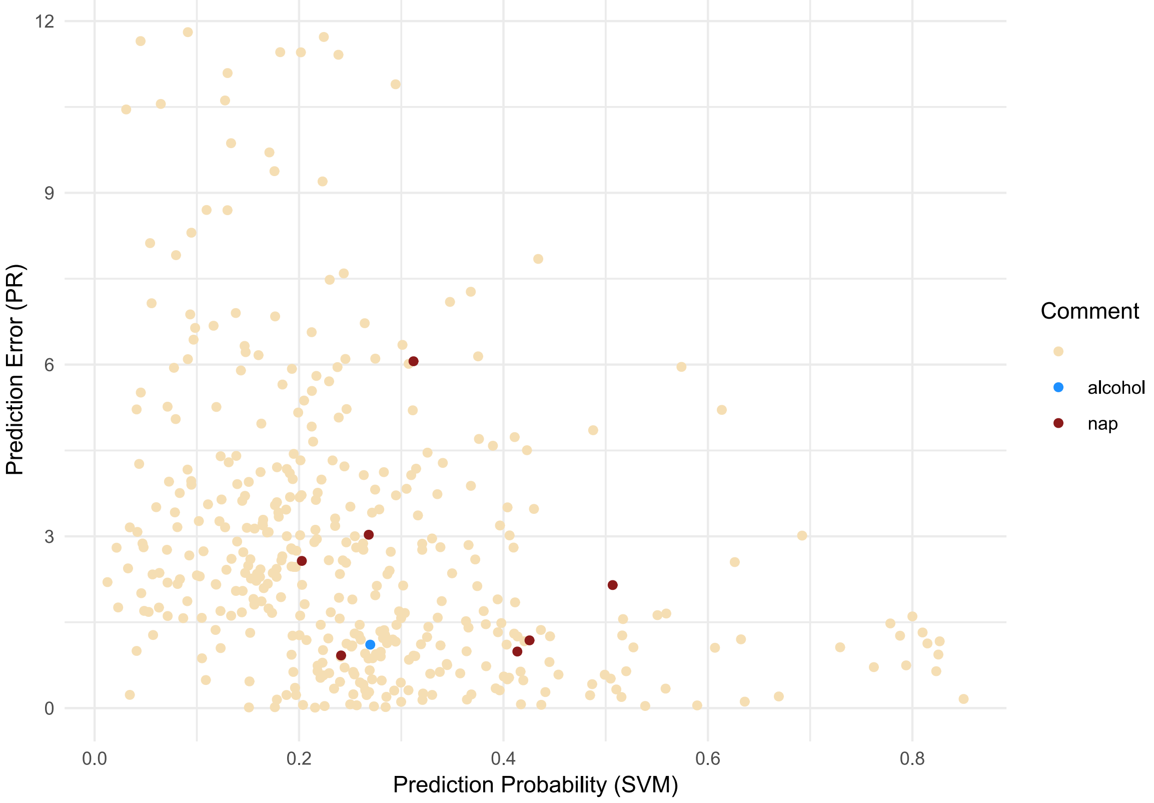


**Suppl. Figure 2: Prediction error and predicted probability shown in relation to donor characteristics and activities:**
Predictions obtained for samples in the respective test set of five-fold cross-validation (n=408 samples from 51 individuals deposited at eight different timepoints, dried for 48 hours; Refall normalised dataset)
**a:** Donor sex, **b:** Donor age (categorical) , **c:** Donor chronotype (assessed using the D-MEQ questionnaire), **d:** Time of the year of sample deposition, **e:** Consumption of caffeine (within 3 hours prior to sample deposition) , **f:** Deviations from study protocol instructions (consumption of alcohol or napping within 3 hours prior to sample deposition) , **g:** Individuals taking medication (medication indicated in brackets (vitamins and oral contraceptives not included)).

**g** Relationship between absolute error and prediction probability

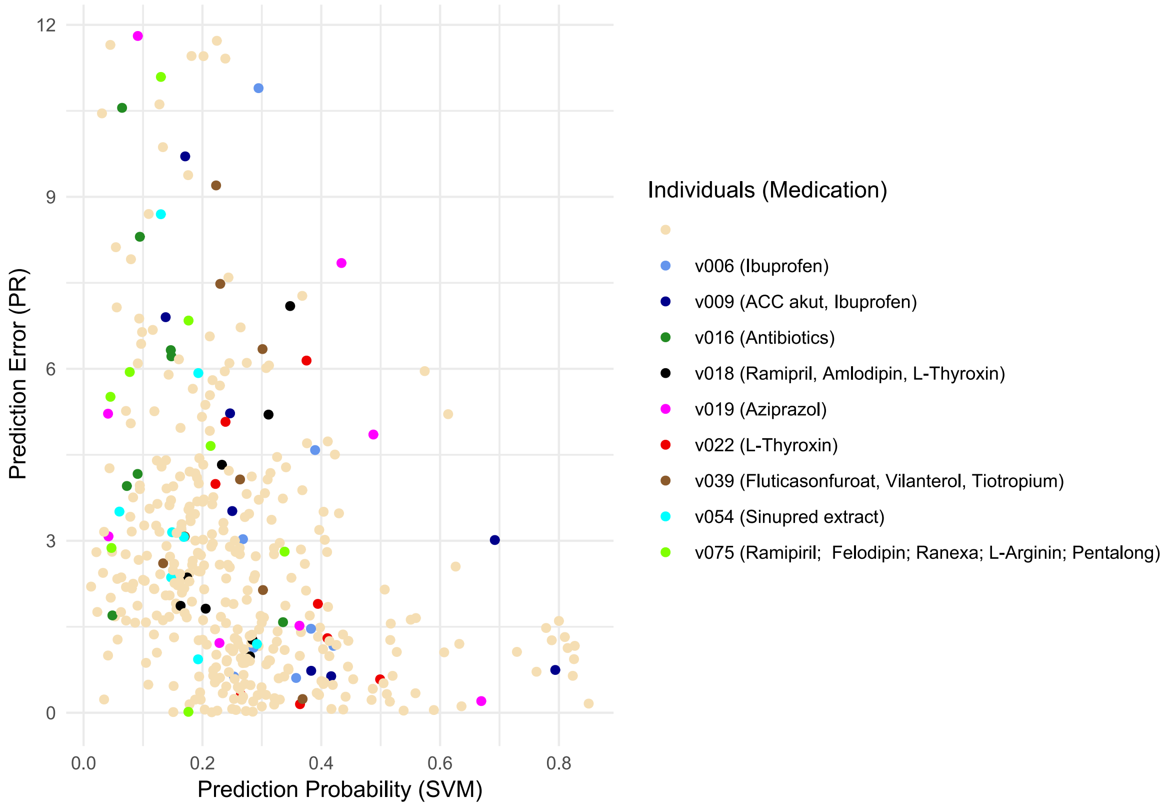


**Suppl. Figure 2: Prediction error and predicted probability shown in relation to donor characteristics and activities:**
Predictions obtained for samples in the respective test set of five-fold cross-validation (n=408  samples from 51 individuals deposited at eight different timepoints, dried for 48 hours; Refall normalised dataset)
**a:** Donor sex, **b:** Donor age (categorical) , **c:** Donor chronotype (assessed using the D-MEQ questionnaire), **d:** Time of the year of sample deposition, **e:** Consumption of caffeine (within 3 hours prior to sample deposition) , **f:** Deviations from study protocol instructions (consumption of alcohol or napping within 3 hours prior to sample deposition) , **g:** Individuals taking medication (medication indicated in brackets (vitamins and oral contraceptives not included)).

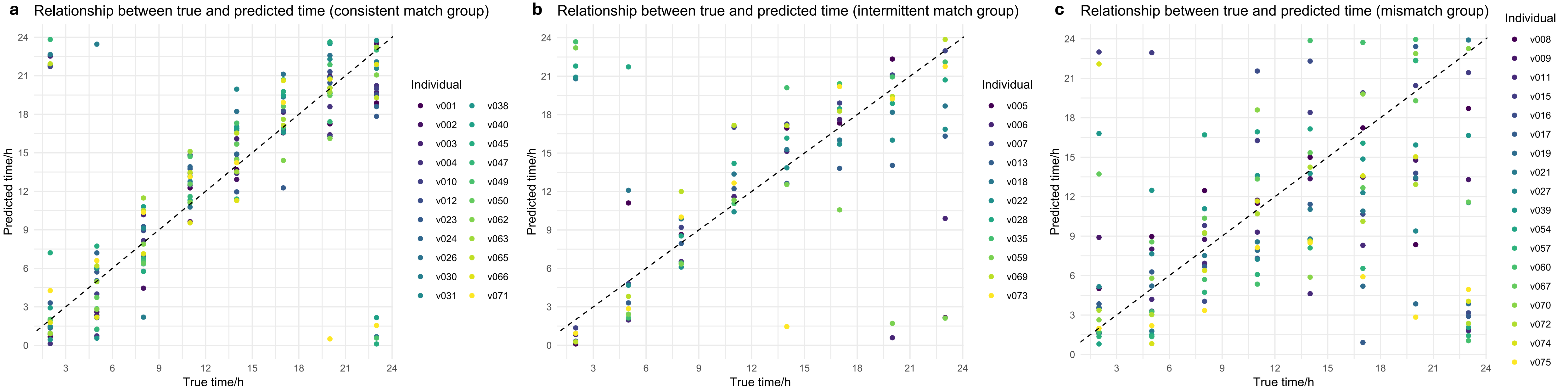


**Suppl. Figure 3: Scatterplot showing true and predicted times for individuals according to match group**

Predictions obtained for samples in the respective test set of five-fold cross-validation (n=408 samples from 51 individuals deposited at eight different timepoints, dried for 48 hours; Refall normalised dataset)

**a:** consistent match group (individuals showing no prediction error >6 hours and a maximum of four samples with a prediction error > 3 hours), **b:** intermittent match group (individuals with a single sample showing a prediction error > 6 hours and a maximum of three samples with a prediction error > 3 hours), **c:** mismatch group (individuals showing a single sample with a prediction error > 6 hours and a minimum of four samples with an error > 3 hours or two or more samples with a prediction error > 6 hours).

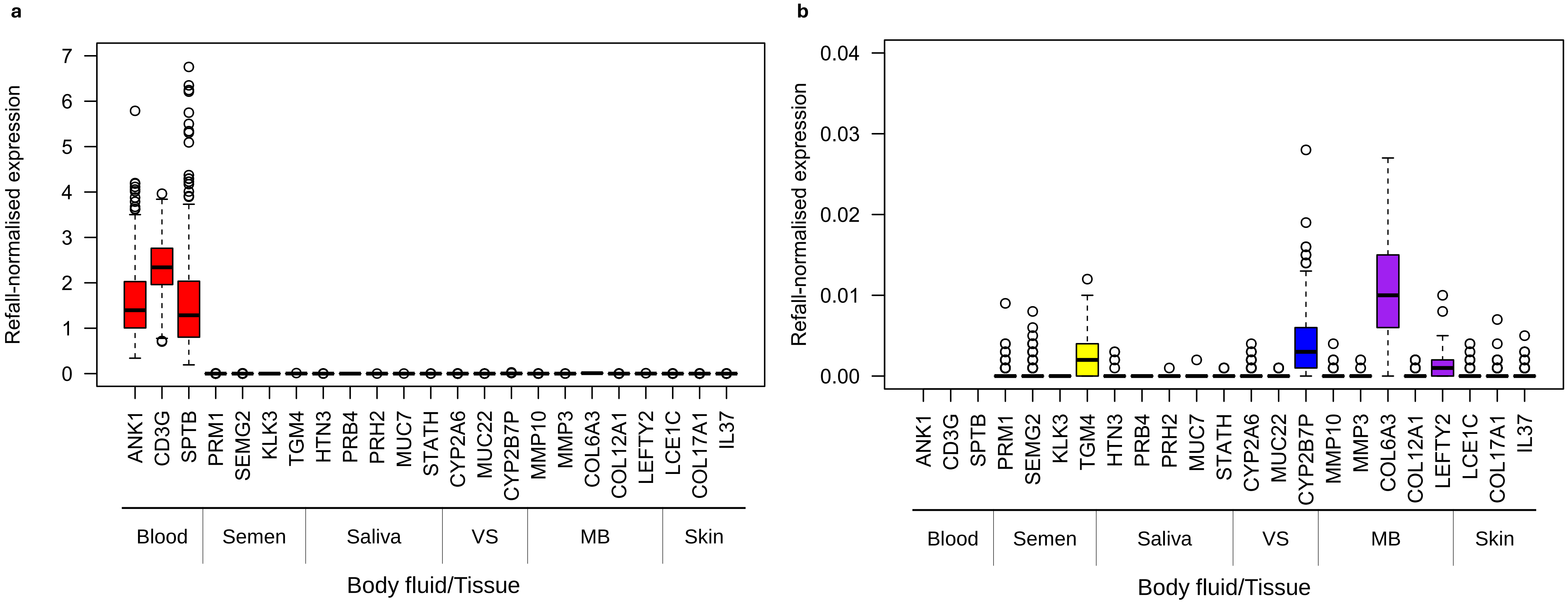


**Suppl. Figure 4: Boxplot showing Refall-normalised expression of BFI markers in blood samples**

Boxplots show the distribution of average values (calculated from n = 408 samples from 51 different individuals collected at 8 different timepoints) for 23 BFI markers. Boxes indicate median and interquartile range (IQR) of this distribution, whiskers extend from the IQR to the largest/smallest value no further than 1.5x IQR. Data points beyond whiskers are plotted individually. Boxes for markers specific for the same body fluid are shown in the same color.

**a:** Plot with y-axis spanning from 0 to 7, **b:** Plot with y-axis spanning from 0 to 0.04 (for better visualisation of low-expressed markers).

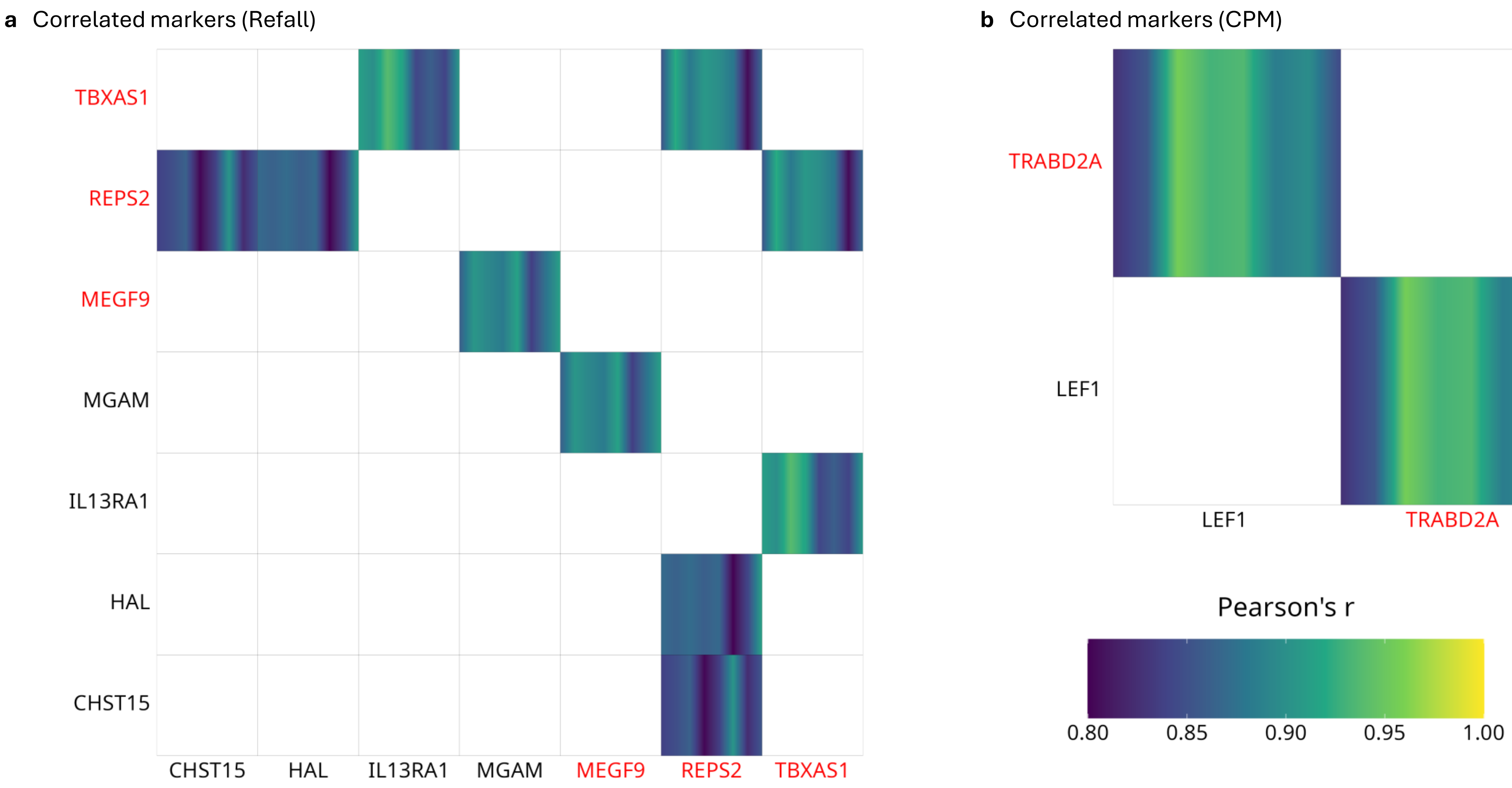


**Suppl. Figure 5: Correlation and selection of markers**
Those markersare shown where all expressions of the 8 time points show a Pearson correlation |r| > 0.8 with those of another marker at the respective time point. The colour gradient marks the correlation accross the eight timepoints from left to right. The genes for which no correlations above the threshold remain after removal are marked in red. **a:** Refall-normalised dataset, **b:** CPM-normalised dataset.
