## Supplementary File 1 for "TrACES of Time: Towards estimating time-of-day of bloodstain deposition by targeted RNA sequencing"

**Supplementary File 1: Comparison of normalisation procedures for targeted RNA sequencing data**

### 1 – Normalisation procedures

Gene expression quantification methods require normalisation procedures to account for technical variation in the various sample preparation steps. While robust normalisation procedures have been established and guidelines have been proposed for the analysis of quantitative PCR data [1, 2], microarray experiments [3] as well as whole transcriptome sequencing approaches [4], no such standards or guidelines have been established for the analysis of data from targeted sequencing of cDNA.

We thus chose to apply and compare six different normalisation procedures (which have been applied in previous studies using targeted RNA sequencing or adapted from other technical approaches such as qPCR or whole transcriptome sequencing), summarised in SF1-Table 1.

**SF1-Table 1: Description of normalisation procedures**

| Name/abbreviation of procedure | Description |
|---|---|
| "CPM" (Counts Per Million) | For each sample, the raw read count of each target gene was divided by the total read counts of all target genes and multiplied by one million. CPM thus represents the number of counts expected to be observed for the respective target gene if each library was sequenced to a total of exactly one million reads. |
| "Median" | For each sample, the raw read count of each target gene was divided by the median of the read counts of all targeted genes. |
| "Refall" | For each sample, the raw read count of each target gene was divided by the average read counts of all reference gene candidates (*DECR1, FPGS, HGS, KIDINS220, MRFAP1, OTDU5, PPIB, TRAP1, USP19*) in that same sample. |
| "Ref3-q" | For each sample, the raw read count of each target gene was divided by the average read counts of a subset of three reference gene candidates (*HGS, KIDINS220, PPIB*) in that same sample. These genes were chosen as they had been identified as the most stable candidates in the qPCR dataset in [5]. |
| "Ref3-NR" | For each sample, the raw read count of each target gene was divided by the average read counts of a subset of three reference gene candidates (*HGS, KIDINS220, TRAP1*) in that same sample. These genes were chosen as they had been identified as the most stable combination of three reference gene candidates when applying NormiRazor [6] to our targeted RNA sequencing dataset. |
| "DEseq2-norm"[1] | This normalisation approach is implemented in the DEseq2 [7] R package and regularly applied to normalise whole transcriptome sequencing data.<br>Briefly summarised, the function creates a 'pseudo-reference sample', to which, for each gene, the geometric mean across all samples is assigned as 'pseudo-read count'. For each gene in each sample, the ratio of the measured read count to this 'pseudo-read count is calculated. All read counts in each sample are then scaled by a size factor, which is set to the median of the calculated ratios per sample [8]. |

[1]Note: The normalisation approach implemented in the DEseq2 package is typically applied to whole transcriptome sequencing data and normalises over the entire dataset. It will therefore change if different samples are combined for normalisation. While this is unproblematic for whole transcriptome sequencing data, where biostatistical analyses are commonly performed on a fixed sample set, this would be an issue for the application envisioned here, where biostatistical models built on a fixed dataset are meant to be applied to single samples sequenced under identical conditions, but possibly much later. Hence, the DEseq2-norm approach and similar approaches are not well suited for our application, however, the DEseq2-norm approach was included in our initial testing for comparison purposes. All other normalisation procedures can be performed on raw data from individual samples and are thus dataset-independent.

**2 – Choice of reference genes**

The reference gene candidates were chosen as described in [5], combining markers identified as stably expressed in human blood in previous studies [9, 10] as well as in our own whole transcriptome sequencing dataset [11]. Despite that, we observed that individual reference genes showed timepoint-dependent expression (Suppl. Table 7). However, as discussed by Kirk et al. [12], normalisation to reference genes with steady expression is strictly necessary only if the aim of the study is to accurately quantify the effect of a certain condition on gene expression, whereas for the aim of identifying patterns in gene expression suitable to predict time-of-day, even markers with cyclic expression patterns might be suitable for normalisation. Additionally, the above-described reference gene candidates were also chosen for their previously described expression stability across individuals and therefore may still be deemed suitable to normalise for interindividual differences in gene expression. We thus decided to include the reference gene-based normalisation approaches in our comparisons despite timepoint dependence of individual reference markers observed during initial marker suitability evaluation.

**3 – Qualitative comparison of normalisation procedures**

The comparison of normalisation procedures was performed on a subset of samples comprising a total of 180 samples from 45 individuals taken at four different time points (8 AM, 2 PM, 8 PM and 2 AM, all stains dried and stored for 48 hours). Raw UMI counts from all 180 samples were normalised applying the six approaches described in SF1-Table 1.

In a first step, we aimed to visually inspect general trends in the datasets. For this purpose, Principal Component Analysis (PCA) of each normalised dataset was performed and PCA plots generated using the "stats" (v. 4.1.1) and the "factoextra" (v. 1.0.7) packages in R [13, 14]. When considering all 69 potential time-of-day predictor candidates, clustering of datapoints according to time-of-day of sample collection was not observed in any of the normalised datasets (SF1-Figure 1).

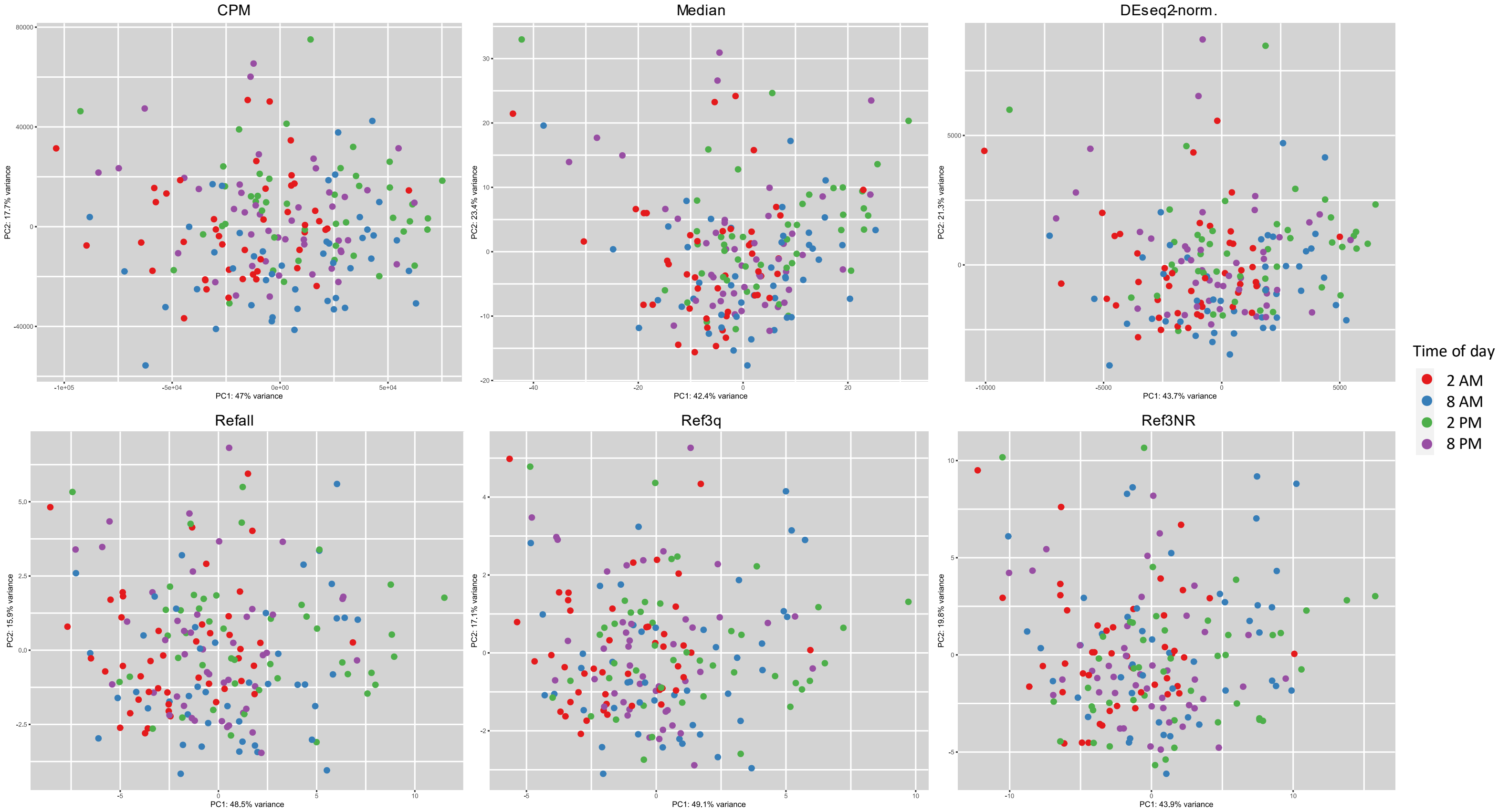


**SF1-Figure 1: PCA plots of normalised gene expression data (all markers)**
Raw read counts were normalised according to the 6 procedures described in SF1-Table 1. The PCA was calculated for all 69 "time-of-day" candidate markers for a subset of 180 samples from 45 individuals taken at four different time points (stains dried for 48 hours). Colours indicate time-of-day of sample deposition.

However, initial data analysis indicated that not all 69 time-of-day predictor candidates were indeed strongly correlated with the time-of-day of sample collection in our dataset. A second PCA was thus performed, considering only a subset of candidate predictors (CAMKK1, FAAH2, IGF2R, FKBP5, AVIL, ERMN, IL13RA1; SF1-Figure 2), which had shown promising results in an initial phase of data analysis (data not shown).

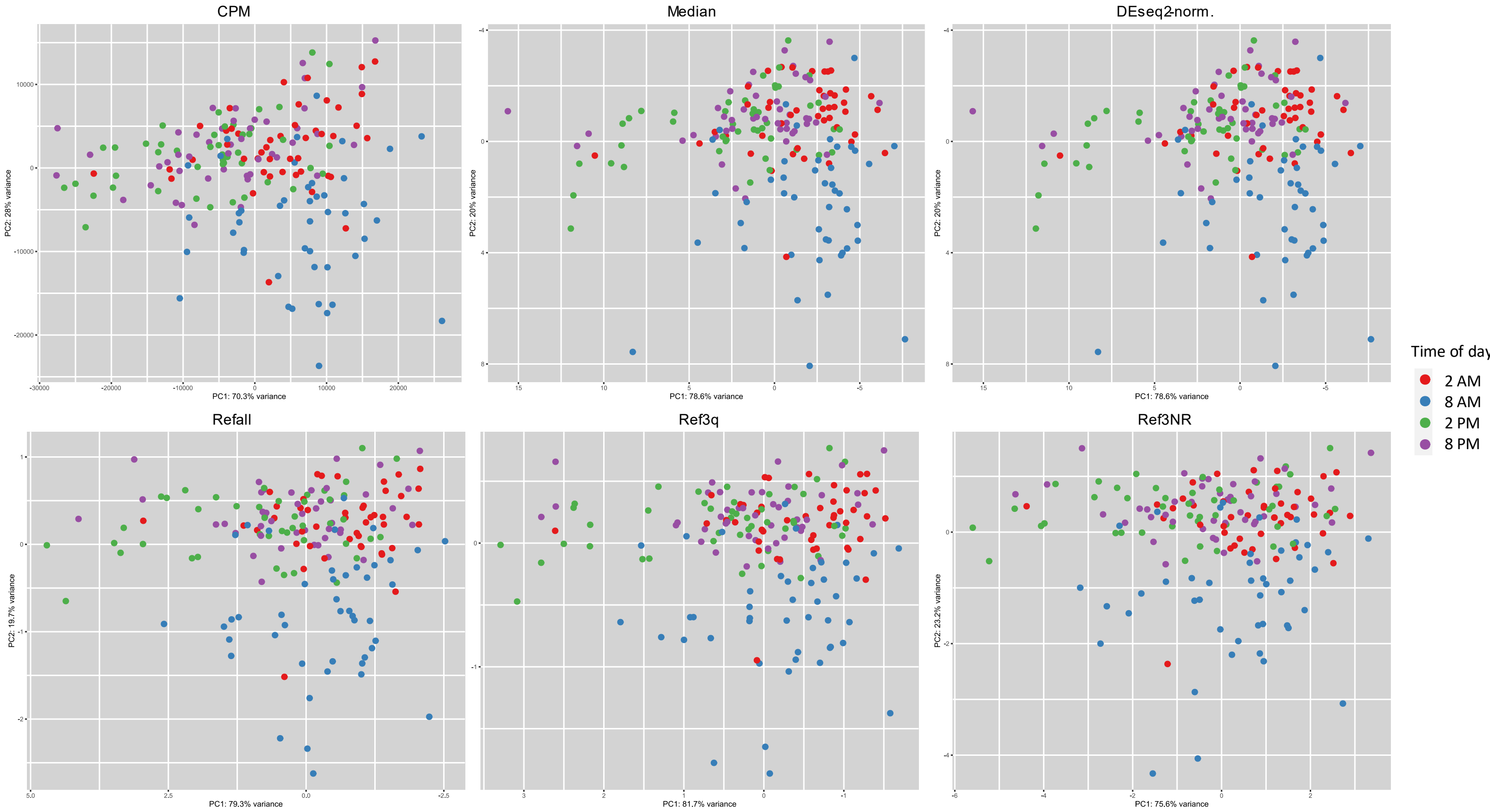


**SF1-Figure 2: PCA plots of normalised gene expression data (selected markers)**
Raw read counts were normalised according to the 6 procedures described in SF1-Table 1. The PCA was calculated for all seven "time-of-day" candidate markers (CAMKK1, FAAH2, IGF2R, FKBP5, AVIL, ERMN, IL13RA1), which had shown promising results in an initial phase of data analysis (data not shown). Analysis was performed for a subset of 180 samples from 45 individuals taken at four different time points (stains dried for 48 hours). Colours indicate time-of-day of sample deposition.

The PCA plots now showed a certain extent of clustering of datapoints according to time-of-day of sample collection, with samples taken at 8 AM showing a more distinct separation than the other groups. Notably, clustering trends were highly comparable between normalisation procedures. Furthermore, all normalised datasets showed a similar proportion of variance explained by the first two principal components. PCA thus did not reveal major differences between the six normalisation procedures.

### 4 – Quantitative comparison of normalisation procedures

In a second step, we aimed to compare the effectiveness of the different normalisation approaches in a quantitative manner. An effective normalisation should remove or at least reduce differences in read counts from technical variability but preserve differences due to true biological differences in gene expression [4]. The "effectiveness" of a normalisation may be assessed in different ways. In this study, we chose to define an effective normalisation procedure as one that reduces intra-group variability (i.e. variability between samples deposited at the same timepoint), while at the same time preserving or even increasing variability between groups and hence used the generalised $\eta^2$ as a measure of normalisation effectiveness.

For this purpose, ANOVAs were calculated using the "rstatix" (v. 0.7.2) package in R [13, 15] and generalised $\eta^2$ values obtained for each gene in each of the six normalised as well as in the unnormalised dataset from the ANOVA table.

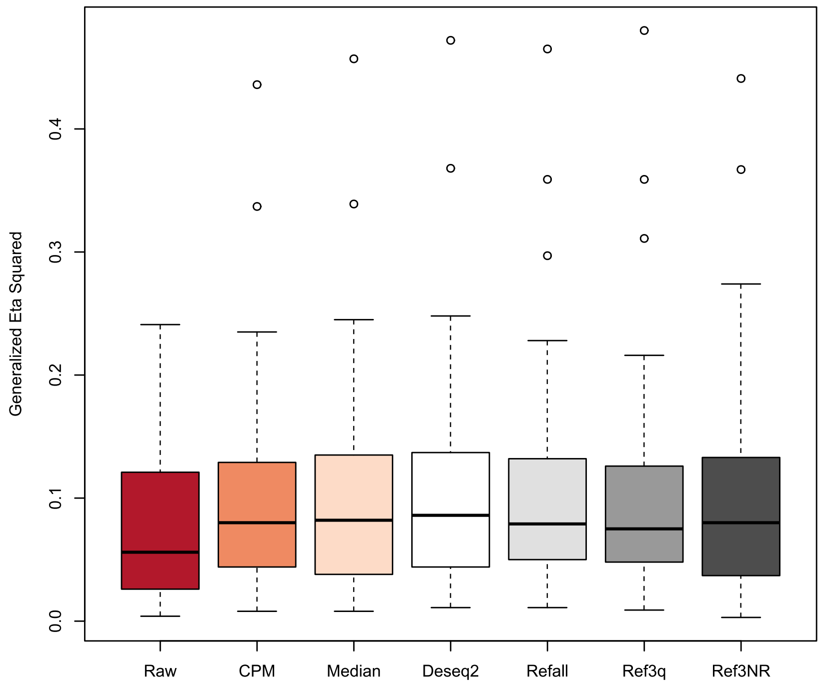


**SF1-Figure 3:**
Raw read counts were normalised according to the six procedures described in SF1-Table 1. Boxplots show the distribution of generalised $\eta^2$ for normalised and raw read counts. Boxes indicate median and interquartile range (IQR) of this distribution, whiskers extend from the IQR to the largest/smallest value no further than 1.5x IQR. Data points beyond whiskers are plotted individually.

SF1-Figure 3 shows the distribution of generalised $\eta^2$ for normalised and raw read counts. Generalised $\eta^2$ increased in all normalised datasets compared to the non-normalised data. It may thus be concluded that each of the six considered approaches is generally suitable for normalisation. While some differences were observed for individual candidate predictors, no normalisation procedure performed vastly better than the others.

**5 – Conclusion**
In consequence, we decided to perform all downstream analyses using two different normalisation approaches from the two conceptionally different procedure groups (normalisation based on reference genes and normalisation based on read count distribution). Within each group, we chose the approach leading to the highest average generalised $\eta^2$ values: Refall and CPM. (The normalisation procedure implemented in the DEseq2 package was considered unsuitable due to the limitation described above. However, our comparisons showed that the read counts normalised with this approach did not differ relevantly from read counts normalised with any of the other approaches).
Considering the results presented in SF1-Figure 1 to 3, it is to be expected that downstream analyses would lead to comparable results if any of the other normalisation procedures would be chosen instead.

**SF1 - References**

1. Bustin SA, Benes V, Garson JA, Hellemans J, Huggett J, Kubista M, et al. The MIQE guidelines: Minimum information for publication of quantitative real-time PCR experiments. Clin Chem. 2009;55:611–22. doi:10.1373/clinchem.2008.112797.
2. Huggett JF, Foy CA, Benes V, Emslie K, Garson JA, Haynes R, et al. The digital MIQE guidelines: Minimum Information for Publication of Quantitative Digital PCR Experiments. Clin Chem. 2013;59:892–902. doi:10.1373/clinchem.2013.206375.
3. Brazma A, Hingamp P, Quackenbush J, Sherlock G, Spellman P, Stoeckert C, et al. Minimum information about a microarray experiment (MIAME)—toward standards for microarray data. Nat Genet. 2001;29:365–71. doi:10.1038/ng1201-365.
4. Evans C, Hardin J, Stoebel DM. Selecting between-sample RNA-Seq normalization methods from the perspective of their assumptions. Brief Bioinformatics. 2018;19:776–92. doi:10.1093/bib/bbx008.
5. Gosch A, Bhardwaj A, Courts C. TrACES of Time: Transcriptomic Analyses for the Contextualization of Evidential Stains – identification of RNA markers for estimating time-of-day of bloodstain deposition. Forensic Science International: Genetics. 2023:102915. doi:10.1016/j.fsigen.2023.102915.
6. Grabia S, Smyczynska U, Pagacz K, Fendler W. NormiRazor: tool applying GPU-accelerated computing for determination of internal references in microRNA transcription studies. BMC Bioinformatics. 2020;21:425. doi:10.1186/s12859-020-03743-8.
7. Love MI, Huber W, Anders S. Moderated estimation of fold change and dispersion for RNA-seq data with DESeq2. Genome Biol. 2014;15:550. doi:10.1186/s13059-014-0550-8.
8. members of the teaching team at the Harvard Chan Bioinformatics Core (HBC). Introduction to DGE. https://hbctraining.github.io/DGE_workshop/lessons/02_DGE_count_normalization.html. Accessed 24 Jun 2024.
9. Hounkpe BW, Chenou F, Lima F de, Paula EV de. HRT Atlas v1.0 database: redefining human and mouse housekeeping genes and candidate reference transcripts by mining massive RNA-seq datasets. Nucleic Acids Res. 2021;49:D947-D955. doi:10.1093/nar/gkaa609.
10. Stamova BS, Apperson M, Walker WL, Tian Y, Xu H, Adamczy P, et al. Identification and validation of suitable endogenous reference genes for gene expression studies in human peripheral blood. BMC Med Genomics. 2009;2:49. doi:10.1186/1755-8794-2-49.
11. Dos Santos KCG, Desgagné-Penix I, Germain H. Custom selected reference genes outperform pre-defined reference genes in transcriptomic analysis. BMC Genomics. 2020;21:35. doi:10.1186/s12864-019-6426-2.
12. Kirk D, Allen R. Exploration of Rhythmic Patterns of Gene Expression to Estimate the Time of Day a Bloodstain Was Created. Research and Reports in Forensic Medical Science. 2021:1–11. doi:10.2147/RRFMS.S327044.
13. R Core Team. R: A Language and Environment for Statistical Computing. Vienna, Austria.
14. Kassambara A, Mundt F. factoextra: Extract and Visualize the Results of Multivariate. 2020. https://CRAN.R-project.org/package=factoextra.
15. Kassambara A. rstatix: Pipe-Friendly Framework for Basic Statistical Tests. https://rpkgs.datanovia.com/rstatix/.
