## Supplementary File 2 for "TrACES of Time: Towards estimating time-of-day of bloodstain deposition by targeted RNA sequencing"

**Supplementary File 2: Optimisation of sequencing depth**

When sequencing libraries are prepared with Unique Molecular Identifiers (UMIs), sufficient sequencing depth is necessary to accurately quantify the number of different UMIs per target marker present in each sample. For sequencing libraries without UMIs, an increase in sequencing reads per sample results in a larger number of reads per marker (thus, accounting for sequencing depth is strictly necessary for read count normalisation). For UMI-based sequencing libraries, in theory, an increase in sequencing reads only leads to an increase in UMI reads as long as not all unique UMI sequences have been sequenced yet. After this point, further increasing sequencing depth will result in an increase in reads per UMI while the UMI read counts will reach a plateau. The sequencing depth at which this plateau is reached will differ depending on the target gene expression level in the individual sample as well as the expression levels of all other markers in the same sample and thus needs to be individually determined for each marker panel and sample type. As targeted cDNA sequencing especially using UMI-based protocols is not widely used for gene expression quantification yet, gold standard procedures or guidelines how to set a minimum sequencing depth threshold are not available to this point.

For the purpose of this study, we aimed to identify the sequencing depth at which the plateau is reached for the majority of markers by performing rarefaction analysis, i.e. starting at a certain sequencing depth and downsampling to different percentages of reads. For rarefaction analysis, we selected eight samples from eight different individuals (two samples per deposition time point 8 AM, 2 PM, 8 PM and 2 AM), sequenced at (or downsampled to) a starting depth of ~600.000 reads per sample. The starting depth was chosen as representing approximately 1.4x the depth recommended by the manufacturer (recommendation in the manufacturer's manual: 5,000 reads per marker for medium sequencing depth; resulting in a recommended sequencing depth of 430,000 reads per sample for a marker panel for which expression of 86 markers is expected (no expression is expected for gDNA controls and non-blood body fluid markers). Reads were then downsampled to 90%, 80%, 70%, 60%, 60%, 25% and 10% of the original 600.000 reads per sample. Downsampling of basecaller.bam files was performed using the filterBam function from the Rsamtools R package (v. 2.18.0) [1]. The UMI read counts per marker and the average reads per UMI for each marker in each downsampled file were compared to the original read file (~600,000 reads).

For two markers (*ATG2A* and *IRS2*), an almost linear increase in UMI read counts was observed with increased sequencing depth. For these markers, increasing the sequencing depth did not at all (*ATG2A*) or only slightly (*IRS2*) lead to an increase in the average number of reads per UMI. This was interpreted as an indication of some malfunction in the primer binding or amplification process and these two markers were thus excluded from further analysis.

The reduction of raw and Refall-normalised UMI read counts per marker and average reads per UMI for the remaining 81 time of day candidate prediction markers is shown in SF2-Figure 1. Per marker values are presented in SF2-Tables 1-3 at the end of this file.

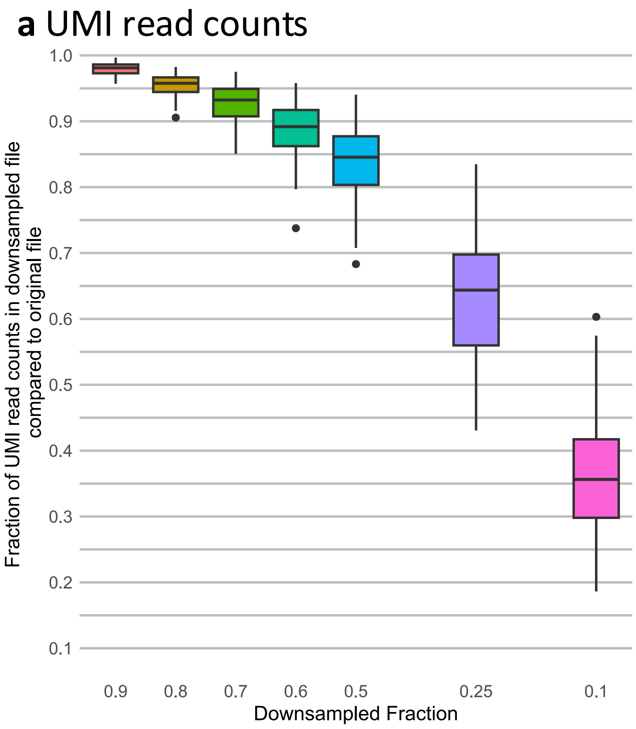


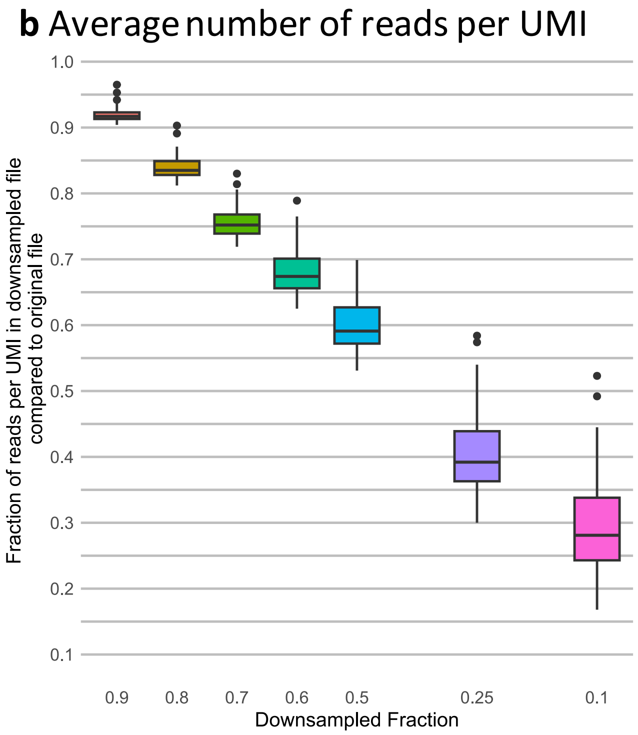


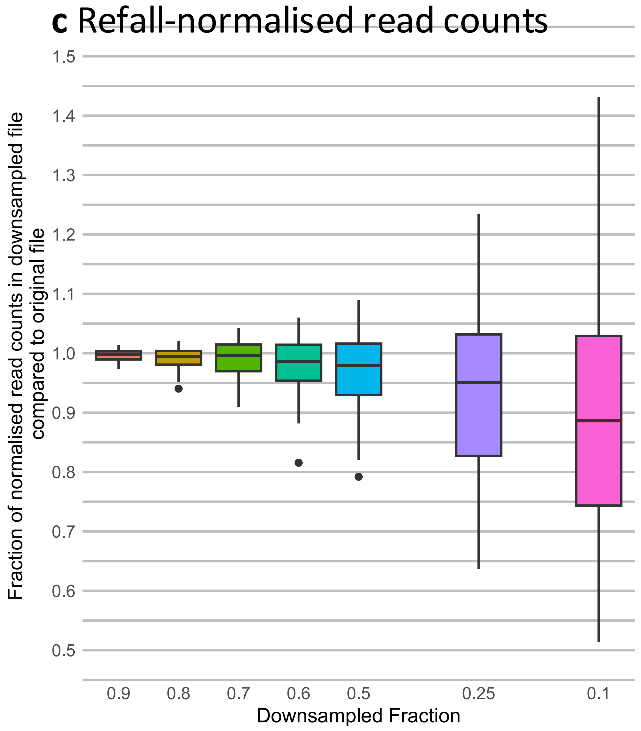


**SF2-Figure 1: Rarefaction analysis**

**a** Comparison of UMI read counts in downsampled fraction compared to original file (~600,000 reads) **b** Comparison of the average number of reads per UMI in downsampled fraction compared to original file (~600,000 reads) **c** Comparison of Refall-normalised read-counts in downsampled fraction compared to original file (~600,000 reads); boxplots show the distribution of average values (calculated from n = 8 samples from eight different individuals (two samples per deposition time point 8 AM, 2 PM, 8 PM and 2 AM)) for 81 candidate time of day prediction markers (excluding IRS2 and ATG2A). Boxes indicate median and interquartile range (IQR) of this distribution, whiskers extend from the IQR to the largest/smallest value no further than 1.5x IQR. Data points beyond whiskers are plotted individually. Box colours indicate the downsampled fraction.

SF2-Figure 1 shows that an increase in sequencing depths only leads to a small increase in UMI read counts for the highest downsampling steps while an almost linear increase in average number of reads per UMI can be observed. The variation in normalised read counts in the downsampled fractions is small compared to the original file for the highest downsampling steps (SF2-Figure 1c). This indicates saturation or near-saturation was achieved for the majority of markers. Further increasing the sequencing depth would lead to a further increase of reads per UMI but no relevant increase in UMI read counts is to be expected.

Based on SF2-Figure 1 we conclude that saturation was achieved for the majority of markers when downsampled to 80% (corresponding to a sequencing depth of ~480,000 reads per sample (≈400,000 reads used for UMI counting)). We thus decided to aim for a sequencing depth resulting in a minimum of ~400,000 reads used for UMI counting per sample.

To evaluate the validity of our approach, we assessed the impact of sequencing depth on prediction error in our final model:

Visual inspection does not indicate a direct relationship between the number of reads used for UMI counting and prediction error (SF2-Figure 2).

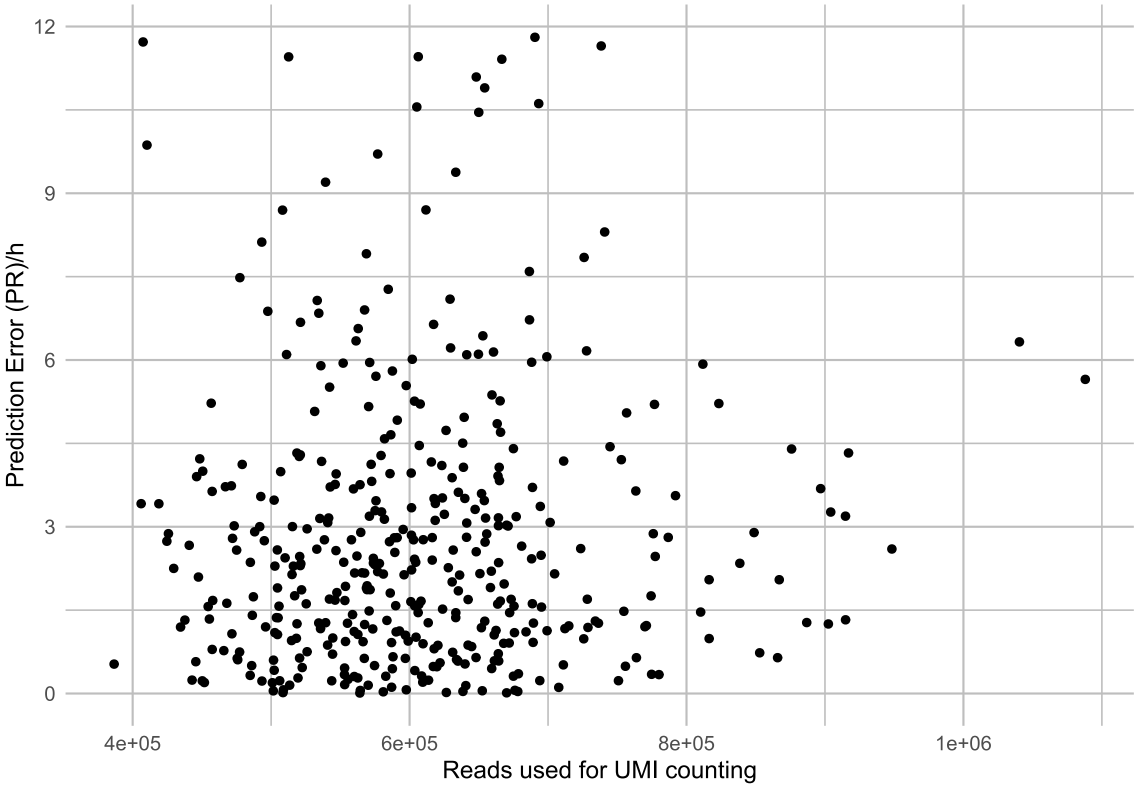


**SF2-Figure 2: Relationship between absolute error and sequencing depth**
Plot showing the relationship between the number of reads used for UMI counting and the absolute prediction error obtained for samples in the respective test set in five-fold cross-validation for the final model (Penalised Regression on Refall-normalised dataset, n=408 dried blood samples from 51 individuals deposited at eight different timepoints).

Additionally, the impact of sequencing depth on prediction error was assessed by including the number of reads used for UMI counting in a mixed-effects model as described in section 5.3.9. In accordance with SF2-Figure 2, the prediction error was not significantly impacted by the number of reads used for UMI counting ($p_{LMM} > 0.05$).

Finally, we evaluated whether excluding the markers showing the highest risk of not having fully reached saturation at the chosen minimum sequencing depth would improve prediction performance. These markers were identified as markers showing an average deviation of >3% in the 80% downsampled fraction compared to the original file or a deviation of >5% for at least one of the eight samples. According to these criteria, 18 markers were identified as potentially not having fully reached saturation (highlighted in orange in SF2-Table 3). Of these, seven were amongst the ten highest-ranking markers in the sine or the cosine component of the final model (Table 2). We therefore excluded these seven markers and constructed a new PR model on the Refall-normalised dataset described in sections 5.3.4 and 5.3.5. Using five-fold cross-validation, we obtained an RMSE of 3.89 h (a mean absolute prediction error of 2.92 h, and a prediction accuracy within +/- 4 hours of 0.77), hence representing a small decrease in prediction accuracy compared to the original model including these seven markers (RMSE of 3.73 h, mean absolute prediction error of 2.81 h and a prediction accuracy within +/- 4 hours of 0.78, cf. Table 1).

From these results, we conclude that the sequencing depth used in this study was sufficient for the sample type and marker panel analysed here. In addition, we note the necessity of developing gold standard procedures and guidelines for this type of analysis to ensure validity and comparability of results in future studies.

**SF2-Table 1: Rarefaction analysis – UMI read counts**

Comparison of UMI read counts in downsampled fraction compared to original file (~600,000 reads). The table shows average values calculated from n = 8 samples from eight different individuals (two samples per deposition time point 8 AM, 2 PM, 8 PM and 2 AM)) for 72 candidate time of day prediction markers and 9 references genes.

| | Downsampled Fraction | | | | | | |
|---|---|---|---|---|---|---|---|
| **Marker** | 0.9 | 0.8 | 0.7 | 0.6 | 0.5 | 0.25 | 0.1 |
| **ABCD2** | 0.990 | 0.969 | 0.937 | 0.934 | 0.869 | 0.703 | 0.429 |
| **ABHD5** | 0.980 | 0.956 | 0.927 | 0.889 | 0.844 | 0.630 | 0.352 |
| **ALOX5AP** | 0.979 | 0.955 | 0.920 | 0.885 | 0.833 | 0.627 | 0.342 |
| **APMAP** | 0.970 | 0.940 | 0.902 | 0.851 | 0.793 | 0.545 | 0.280 |
| **ARG1** | 0.986 | 0.962 | 0.951 | 0.907 | 0.877 | 0.676 | 0.418 |
| **AVIL** | 0.986 | 0.957 | 0.939 | 0.894 | 0.859 | 0.680 | 0.399 |
| **BACH2** | 0.983 | 0.960 | 0.932 | 0.891 | 0.850 | 0.655 | 0.366 |
| **BCL2** | 0.987 | 0.972 | 0.954 | 0.931 | 0.890 | 0.738 | 0.464 |
| **CAMKK1** | 0.960 | 0.920 | 0.885 | 0.833 | 0.733 | 0.472 | 0.223 |
| **CHST15** | 0.967 | 0.928 | 0.881 | 0.819 | 0.758 | 0.508 | 0.250 |
| **CLEC4E** | 0.984 | 0.965 | 0.942 | 0.914 | 0.874 | 0.685 | 0.397 |
| **CLIC3** | 0.984 | 0.944 | 0.910 | 0.869 | 0.815 | 0.596 | 0.320 |
| **CNTLN** | 0.986 | 0.964 | 0.942 | 0.908 | 0.873 | 0.676 | 0.400 |
| **CPD** | 0.981 | 0.955 | 0.922 | 0.883 | 0.824 | 0.610 | 0.340 |
| **CRISPLD2** | 0.968 | 0.950 | 0.908 | 0.860 | 0.783 | 0.560 | 0.275 |
| **DAAM2** | 0.997 | 0.941 | 0.958 | 0.880 | 0.856 | 0.712 | 0.401 |
| **DECR1** | 0.975 | 0.950 | 0.916 | 0.871 | 0.810 | 0.590 | 0.306 |
| **ECHDC3** | 0.986 | 0.965 | 0.958 | 0.927 | 0.903 | 0.725 | 0.412 |
| **ENHO** | 0.984 | 0.975 | 0.926 | 0.890 | 0.803 | 0.667 | 0.326 |
| **ERGIC1** | 0.969 | 0.932 | 0.892 | 0.836 | 0.776 | 0.530 | 0.262 |
| **ERMN** | 0.984 | 0.961 | 0.938 | 0.897 | 0.857 | 0.680 | 0.400 |
| **FAAH2** | 0.971 | 0.938 | 0.869 | 0.829 | 0.766 | 0.528 | 0.259 |
| **FBXL16** | 0.980 | 0.979 | 0.953 | 0.921 | 0.915 | 0.712 | 0.434 |

| | Downsampled Fraction | | | | | | |
|---|---|---|---|---|---|---|---|
| **Marker** | 0.9 | 0.8 | 0.7 | 0.6 | 0.5 | 0.25 | 0.1 |
| **FKBP5** | 0.982 | 0.959 | 0.933 | 0.904 | 0.858 | 0.662 | 0.382 |
| **FLT3** | 0.972 | 0.945 | 0.957 | 0.916 | 0.882 | 0.690 | 0.442 |
| **FOSL2** | 0.972 | 0.936 | 0.899 | 0.845 | 0.781 | 0.532 | 0.272 |
| **FPGS** | 0.993 | 0.979 | 0.968 | 0.955 | 0.929 | 0.804 | 0.570 |
| **GABARAPL1** | 0.973 | 0.944 | 0.910 | 0.862 | 0.806 | 0.583 | 0.311 |
| **GCNT4** | 0.984 | 0.961 | 0.943 | 0.902 | 0.876 | 0.669 | 0.418 |
| **HAL** | 0.978 | 0.951 | 0.918 | 0.880 | 0.833 | 0.607 | 0.328 |
| **HDAC9** | 0.982 | 0.966 | 0.932 | 0.897 | 0.847 | 0.655 | 0.377 |
| **HGS** | 0.970 | 0.935 | 0.889 | 0.836 | 0.779 | 0.537 | 0.275 |
| **HNRNPDL** | 0.968 | 0.927 | 0.880 | 0.826 | 0.756 | 0.503 | 0.255 |
| **HPCAL4** | 0.980 | 0.978 | 0.937 | 0.877 | 0.819 | 0.625 | 0.314 |
| **ID3** | 0.985 | 0.948 | 0.913 | 0.892 | 0.838 | 0.641 | 0.372 |
| **IGF2R** | 0.974 | 0.940 | 0.906 | 0.854 | 0.793 | 0.546 | 0.283 |
| **IL13RA1** | 0.970 | 0.927 | 0.900 | 0.842 | 0.767 | 0.532 | 0.271 |
| **IL18R1** | 0.973 | 0.931 | 0.882 | 0.815 | 0.776 | 0.521 | 0.260 |
| **IL1R1** | 0.972 | 0.949 | 0.895 | 0.865 | 0.804 | 0.560 | 0.302 |
| **IL1R2** | 0.987 | 0.965 | 0.942 | 0.908 | 0.870 | 0.675 | 0.398 |
| **KIDINS220** | 0.980 | 0.950 | 0.912 | 0.876 | 0.820 | 0.602 | 0.316 |
| **KLF9** | 0.979 | 0.955 | 0.920 | 0.885 | 0.826 | 0.612 | 0.334 |
| **LEF1** | 0.991 | 0.979 | 0.963 | 0.945 | 0.919 | 0.781 | 0.516 |
| **MAL** | 0.981 | 0.959 | 0.930 | 0.895 | 0.845 | 0.644 | 0.356 |
| **MEGF9** | 0.981 | 0.958 | 0.927 | 0.890 | 0.846 | 0.624 | 0.340 |
| **MFGE8** | 0.976 | 0.953 | 0.920 | 0.878 | 0.834 | 0.636 | 0.346 |
| **MGAM** | 0.972 | 0.934 | 0.890 | 0.848 | 0.785 | 0.551 | 0.280 |
| **MKNK2** | 0.988 | 0.973 | 0.955 | 0.933 | 0.901 | 0.747 | 0.481 |
| **MRFAP1** | 0.987 | 0.972 | 0.950 | 0.928 | 0.896 | 0.730 | 0.456 |
| **MSRB2** | 0.987 | 0.966 | 0.937 | 0.892 | 0.852 | 0.652 | 0.397 |
| **MTHFS** | 0.977 | 0.949 | 0.906 | 0.854 | 0.793 | 0.560 | 0.296 |
| **MYO1E** | 0.990 | 0.974 | 0.969 | 0.947 | 0.932 | 0.809 | 0.543 |
| **NELL2** | 0.990 | 0.978 | 0.962 | 0.943 | 0.921 | 0.802 | 0.551 |
| **NR1D1** | 0.979 | 0.960 | 0.944 | 0.908 | 0.858 | 0.655 | 0.382 |
| **NR1D2** | 0.983 | 0.961 | 0.938 | 0.903 | 0.865 | 0.661 | 0.381 |
| **NRCAM** | 0.984 | 0.960 | 0.935 | 0.894 | 0.865 | 0.634 | 0.341 |
| **OTUD5** | 0.989 | 0.972 | 0.952 | 0.920 | 0.893 | 0.724 | 0.447 |
| **PER1** | 0.982 | 0.965 | 0.935 | 0.904 | 0.863 | 0.666 | 0.392 |
| **PER2** | 0.967 | 0.920 | 0.854 | 0.797 | 0.731 | 0.487 | 0.244 |
| **PER3** | 0.973 | 0.928 | 0.874 | 0.870 | 0.756 | 0.552 | 0.283 |
| **PFKFB2** | 0.984 | 0.955 | 0.932 | 0.877 | 0.838 | 0.613 | 0.321 |
| **PHC2** | 0.979 | 0.953 | 0.928 | 0.883 | 0.828 | 0.620 | 0.345 |
| **PPIB** | 0.989 | 0.975 | 0.956 | 0.934 | 0.901 | 0.740 | 0.468 |
| **REPS2** | 0.979 | 0.957 | 0.925 | 0.893 | 0.852 | 0.632 | 0.363 |
| **SAP30** | 0.977 | 0.952 | 0.915 | 0.874 | 0.835 | 0.627 | 0.343 |

| | Downsampled Fraction | | | | | | |
|---|---|---|---|---|---|---|---|
| **Marker** | 0.9 | 0.8 | 0.7 | 0.6 | 0.5 | 0.25 | 0.1 |
| **SEC14L1** | 0.992 | 0.982 | 0.972 | 0.958 | 0.940 | 0.826 | 0.575 |
| **SIPA1L2** | 0.957 | 0.906 | 0.850 | 0.738 | 0.683 | 0.431 | 0.186 |
| **SMAP2** | 0.989 | 0.968 | 0.947 | 0.917 | 0.879 | 0.694 | 0.408 |
| **SPATA6** | 0.961 | 0.916 | 0.858 | 0.800 | 0.708 | 0.461 | 0.207 |
| **SYTL3** | 0.971 | 0.934 | 0.882 | 0.837 | 0.763 | 0.508 | 0.263 |
| **TBXAS1** | 0.990 | 0.982 | 0.969 | 0.955 | 0.940 | 0.835 | 0.603 |
| **TCF7** | 0.986 | 0.964 | 0.943 | 0.910 | 0.875 | 0.682 | 0.400 |
| **TMCC3** | 0.991 | 0.971 | 0.949 | 0.930 | 0.912 | 0.759 | 0.490 |
| **TPST1** | 0.992 | 0.967 | 0.941 | 0.922 | 0.893 | 0.732 | 0.431 |
| **TRABD2A** | 0.984 | 0.964 | 0.944 | 0.918 | 0.877 | 0.698 | 0.417 |
| **TRAP1** | 0.978 | 0.963 | 0.954 | 0.923 | 0.863 | 0.700 | 0.402 |
| **TSC22D3** | 0.986 | 0.969 | 0.951 | 0.925 | 0.889 | 0.733 | 0.464 |
| **UPB1** | 0.960 | 0.976 | 0.975 | 0.894 | 0.829 | 0.687 | 0.328 |
| **USP19** | 0.965 | 0.931 | 0.880 | 0.820 | 0.758 | 0.531 | 0.248 |
| **XKR8** | 0.991 | 0.981 | 0.970 | 0.943 | 0.923 | 0.786 | 0.523 |
| **ZBTB16** | 0.977 | 0.953 | 0.915 | 0.861 | 0.808 | 0.572 | 0.298 |

**SF2-Table 2: Rarefaction analysis – Average reads per UMI**

Comparison of the average number of reads per UMI in downsampled fraction compared to original file (~600,000 reads). The table shows average values calculated from n = 8 samples from eight different individuals (two samples per deposition time point 8 AM, 2 PM, 8 PM and 2 AM)) for 72 candidate time of day prediction markers and 9 references genes.

| | Downsampled Fraction | | | | | | |
|---|---|---|---|---|---|---|---|
| **Marker** | 0.9 | 0.8 | 0.7 | 0.6 | 0.5 | 0.25 | 0.1 |
| **ABCD2** | 0.912 | 0.821 | 0.739 | 0.636 | 0.558 | 0.358 | 0.237 |
| **ABHD5** | 0.914 | 0.828 | 0.755 | 0.674 | 0.594 | 0.403 | 0.292 |
| **ALOX5AP** | 0.921 | 0.838 | 0.761 | 0.679 | 0.597 | 0.404 | 0.298 |
| **APMAP** | 0.924 | 0.848 | 0.778 | 0.713 | 0.639 | 0.459 | 0.370 |
| **ARG1** | 0.916 | 0.831 | 0.749 | 0.672 | 0.572 | 0.366 | 0.243 |
| **AVIL** | 0.913 | 0.819 | 0.749 | 0.667 | 0.579 | 0.371 | 0.249 |
| **BACH2** | 0.917 | 0.832 | 0.744 | 0.675 | 0.591 | 0.394 | 0.273 |
| **BCL2** | 0.904 | 0.816 | 0.733 | 0.649 | 0.551 | 0.334 | 0.216 |
| **CAMKK1** | 0.926 | 0.870 | 0.792 | 0.737 | 0.672 | 0.511 | 0.416 |
| **CHST15** | 0.923 | 0.865 | 0.789 | 0.730 | 0.659 | 0.502 | 0.410 |
| **CLEC4E** | 0.914 | 0.829 | 0.740 | 0.658 | 0.575 | 0.365 | 0.245 |
| **CLIC3** | 0.914 | 0.851 | 0.760 | 0.688 | 0.615 | 0.419 | 0.314 |
| **CNTLN** | 0.912 | 0.830 | 0.746 | 0.659 | 0.573 | 0.363 | 0.254 |
| **CPD** | 0.914 | 0.840 | 0.758 | 0.680 | 0.599 | 0.412 | 0.304 |
| **CRISPLD2** | 0.928 | 0.839 | 0.775 | 0.702 | 0.634 | 0.449 | 0.352 |
| **DAAM2** | 0.926 | 0.860 | 0.749 | 0.672 | 0.586 | 0.305 | 0.308 |
| **DECR1** | 0.929 | 0.845 | 0.763 | 0.694 | 0.625 | 0.430 | 0.326 |
| **ECHDC3** | 0.912 | 0.839 | 0.739 | 0.653 | 0.553 | 0.363 | 0.217 |

| | Downsampled Fraction | | | | | | |
|---|---|---|---|---|---|---|---|
| **Marker** | 0.9 | 0.8 | 0.7 | 0.6 | 0.5 | 0.25 | 0.1 |
| **ENHO** | 0.934 | 0.844 | 0.740 | 0.701 | 0.627 | 0.396 | 0.242 |
| **ERGIC1** | 0.929 | 0.862 | 0.789 | 0.716 | 0.646 | 0.477 | 0.382 |
| **ERMN** | 0.914 | 0.834 | 0.752 | 0.674 | 0.591 | 0.383 | 0.262 |
| **FAAH2** | 0.929 | 0.866 | 0.791 | 0.731 | 0.657 | 0.487 | 0.395 |
| **FBXL16** | 0.925 | 0.816 | 0.732 | 0.656 | 0.559 | 0.359 | 0.229 |
| **FKBP5** | 0.915 | 0.832 | 0.749 | 0.664 | 0.585 | 0.379 | 0.260 |
| **FLT3** | 0.917 | 0.849 | 0.742 | 0.644 | 0.590 | 0.352 | 0.257 |
| **FOSL2** | 0.922 | 0.852 | 0.774 | 0.716 | 0.643 | 0.469 | 0.366 |
| **FPGS** | 0.907 | 0.816 | 0.719 | 0.625 | 0.535 | 0.308 | 0.172 |
| **GABARAPL1** | 0.922 | 0.841 | 0.768 | 0.694 | 0.623 | 0.426 | 0.322 |
| **GCNT4** | 0.912 | 0.830 | 0.741 | 0.662 | 0.571 | 0.362 | 0.247 |
| **HAL** | 0.916 | 0.844 | 0.756 | 0.681 | 0.598 | 0.425 | 0.304 |
| **HDAC9** | 0.914 | 0.829 | 0.751 | 0.671 | 0.591 | 0.378 | 0.262 |
| **HGS** | 0.929 | 0.853 | 0.787 | 0.712 | 0.645 | 0.458 | 0.372 |
| **HNRNPDL** | 0.937 | 0.870 | 0.796 | 0.733 | 0.662 | 0.504 | 0.398 |
| **HPCAL4** | 0.919 | 0.847 | 0.762 | 0.681 | 0.617 | 0.397 | 0.322 |
| **ID3** | 0.911 | 0.840 | 0.763 | 0.692 | 0.594 | 0.390 | 0.261 |
| **IGF2R** | 0.923 | 0.844 | 0.774 | 0.705 | 0.630 | 0.456 | 0.353 |
| **IL13RA1** | 0.923 | 0.852 | 0.785 | 0.721 | 0.649 | 0.475 | 0.384 |
| **IL18R1** | 0.920 | 0.866 | 0.801 | 0.723 | 0.626 | 0.496 | 0.381 |
| **IL1R1** | 0.924 | 0.825 | 0.760 | 0.692 | 0.617 | 0.436 | 0.324 |
| **IL1R2** | 0.916 | 0.830 | 0.748 | 0.665 | 0.573 | 0.373 | 0.252 |
| **KIDINS220** | 0.915 | 0.843 | 0.763 | 0.680 | 0.605 | 0.418 | 0.312 |
| **KLF9** | 0.923 | 0.843 | 0.762 | 0.679 | 0.606 | 0.413 | 0.306 |
| **LEF1** | 0.909 | 0.820 | 0.726 | 0.634 | 0.544 | 0.321 | 0.197 |
| **MAL** | 0.916 | 0.835 | 0.749 | 0.674 | 0.589 | 0.382 | 0.283 |
| **MEGF9** | 0.919 | 0.836 | 0.760 | 0.680 | 0.589 | 0.407 | 0.298 |
| **MFGE8** | 0.925 | 0.845 | 0.760 | 0.672 | 0.599 | 0.398 | 0.295 |
| **MGAM** | 0.922 | 0.851 | 0.787 | 0.707 | 0.636 | 0.459 | 0.361 |
| **MKNK2** | 0.913 | 0.821 | 0.737 | 0.645 | 0.559 | 0.334 | 0.211 |
| **MRFAP1** | 0.913 | 0.826 | 0.740 | 0.648 | 0.561 | 0.347 | 0.220 |
| **MSRB2** | 0.911 | 0.825 | 0.748 | 0.673 | 0.591 | 0.383 | 0.267 |
| **MTHFS** | 0.915 | 0.835 | 0.770 | 0.707 | 0.629 | 0.457 | 0.348 |
| **MYO1E** | 0.911 | 0.816 | 0.723 | 0.633 | 0.534 | 0.318 | 0.188 |
| **NELL2** | 0.909 | 0.820 | 0.727 | 0.638 | 0.542 | 0.311 | 0.184 |
| **NR1D1** | 0.923 | 0.828 | 0.744 | 0.664 | 0.582 | 0.380 | 0.266 |
| **NR1D2** | 0.918 | 0.830 | 0.747 | 0.667 | 0.577 | 0.380 | 0.261 |
| **NRCAM** | 0.910 | 0.832 | 0.754 | 0.678 | 0.580 | 0.366 | 0.274 |
| **OTUD5** | 0.907 | 0.822 | 0.735 | 0.656 | 0.559 | 0.347 | 0.227 |
| **PER1** | 0.917 | 0.830 | 0.751 | 0.669 | 0.577 | 0.379 | 0.257 |
| **PER2** | 0.942 | 0.903 | 0.797 | 0.749 | 0.682 | 0.540 | 0.445 |
| **PER3** | 0.919 | 0.851 | 0.790 | 0.710 | 0.644 | 0.472 | 0.355 |

| | Downsampled Fraction | | | | | | |
|---|---|---|---|---|---|---|---|
| **Marker** | 0.9 | 0.8 | 0.7 | 0.6 | 0.5 | 0.25 | 0.1 |
| **PFKFB2** | 0.916 | 0.852 | 0.761 | 0.683 | 0.602 | 0.418 | 0.309 |
| **PHC2** | 0.924 | 0.851 | 0.754 | 0.683 | 0.606 | 0.406 | 0.288 |
| **PPIB** | 0.908 | 0.820 | 0.732 | 0.640 | 0.561 | 0.339 | 0.219 |
| **REPS2** | 0.917 | 0.835 | 0.758 | 0.677 | 0.583 | 0.392 | 0.281 |
| **SAP30** | 0.921 | 0.832 | 0.760 | 0.681 | 0.598 | 0.405 | 0.299 |
| **SEC14L1** | 0.906 | 0.813 | 0.721 | 0.628 | 0.532 | 0.303 | 0.175 |
| **SIPA1L2** | 0.965 | 0.891 | 0.830 | 0.789 | 0.699 | 0.574 | 0.523 |
| **SMAP2** | 0.911 | 0.828 | 0.743 | 0.654 | 0.567 | 0.366 | 0.246 |
| **SPATA6** | 0.930 | 0.871 | 0.814 | 0.765 | 0.699 | 0.584 | 0.492 |
| **SYTL3** | 0.927 | 0.857 | 0.806 | 0.712 | 0.650 | 0.489 | 0.385 |
| **TBXAS1** | 0.912 | 0.812 | 0.726 | 0.633 | 0.531 | 0.300 | 0.168 |
| **TCF7** | 0.915 | 0.832 | 0.739 | 0.658 | 0.573 | 0.369 | 0.254 |
| **TMCC3** | 0.911 | 0.820 | 0.733 | 0.651 | 0.545 | 0.329 | 0.197 |
| **TPST1** | 0.914 | 0.834 | 0.739 | 0.650 | 0.569 | 0.342 | 0.229 |
| **TRABD2A** | 0.910 | 0.827 | 0.733 | 0.653 | 0.572 | 0.363 | 0.249 |
| **TRAP1** | 0.922 | 0.828 | 0.731 | 0.653 | 0.583 | 0.366 | 0.238 |
| **TSC22D3** | 0.914 | 0.826 | 0.734 | 0.652 | 0.563 | 0.340 | 0.217 |
| **UPB1** | 0.953 | 0.838 | 0.737 | 0.669 | 0.627 | 0.435 | 0.316 |
| **USP19** | 0.931 | 0.858 | 0.793 | 0.740 | 0.671 | 0.484 | 0.388 |
| **XKR8** | 0.908 | 0.819 | 0.726 | 0.634 | 0.541 | 0.315 | 0.188 |
| **ZBTB16** | 0.918 | 0.829 | 0.765 | 0.688 | 0.610 | 0.439 | 0.338 |

**SF2-Table 3: Rarefaction analysis – Refall-normalised read counts**

Comparison of Refall-normalised read-counts in downsampled fraction compared to original file (~600,000 reads). The table shows average values calculated from n = 8 samples from eight different individuals (two samples per deposition time point 8 AM, 2 PM, 8 PM and 2 AM)) for 72 candidate time of day prediction markers and 9 references genes. Markers with the highest risk of not fully having reached saturation (i.e. markers showing an average deviation of >3% in the 80% downsampled fraction compared to the original file or a deviation of >5% for at least one of the eight samples) are highlighted in orange.

| | Downsampled Fraction | | | | | | |
|---|---|---|---|---|---|---|---|
| **Marker** | 0.9 | 0.8 | 0.7 | 0.6 | 0.5 | 0.25 | 0.1 |
| **ABCD2** | 1.007 | 1.006 | 1.002 | 1.033 | 1.007 | 1.039 | 1.067 |
| **ABHD5** | 0.996 | 0.993 | 0.990 | 0.983 | 0.977 | 0.928 | 0.873 |
| **ALOX5AP** | 0.996 | 0.992 | 0.983 | 0.979 | 0.966 | 0.927 | 0.850 |
| **APMAP** | 0.987 | 0.976 | 0.964 | 0.941 | 0.919 | 0.805 | 0.694 |
| **ARG1** | 1.003 | 0.999 | 1.017 | 1.003 | 1.016 | 0.999 | 1.040 |
| **AVIL** | 1.003 | 0.994 | 1.004 | 0.989 | 0.995 | 1.003 | 0.992 |
| **BACH2** | 0.999 | 0.997 | 0.996 | 0.986 | 0.985 | 0.967 | 0.904 |
| **BCL2** | 1.004 | 1.009 | 1.020 | 1.030 | 1.031 | 1.091 | 1.153 |
| **CAMKK1** | 0.976 | 0.956 | 0.946 | 0.921 | 0.848 | 0.697 | 0.556 |
| **CHST15** | 0.983 | 0.964 | 0.942 | 0.906 | 0.878 | 0.750 | 0.620 |

| | Downsampled Fraction | | | | | | |
|---|---|---|---|---|---|---|---|
| **Marker** | 0.9 | 0.8 | 0.7 | 0.6 | 0.5 | 0.25 | 0.1 |
| **CLEC4E** | 1.001 | 1.002 | 1.006 | 1.011 | 1.013 | 1.012 | 0.987 |
| **CLIC3** | 1.001 | 0.980 | 0.972 | 0.962 | 0.944 | 0.884 | 0.802 |
| **CNTLN** | 1.003 | 1.001 | 1.006 | 1.004 | 1.011 | 0.999 | 0.995 |
| **CPD** | 0.998 | 0.991 | 0.985 | 0.977 | 0.954 | 0.901 | 0.845 |
| **CRISPLD2** | 0.984 | 0.986 | 0.970 | 0.951 | 0.908 | 0.827 | 0.684 |
| **DAAM2** | 1.014 | 0.977 | 1.024 | 0.974 | 0.993 | 1.052 | 1.006 |
| **DECR1** | 0.992 | 0.987 | 0.979 | 0.963 | 0.938 | 0.871 | 0.761 |
| **ECHDC3** | 1.003 | 1.002 | 1.024 | 1.025 | 1.047 | 1.071 | 1.023 |
| **ENHO** | 1.001 | 1.013 | 0.990 | 0.986 | 0.930 | 0.986 | 0.774 |
| **ERGIC1** | 0.986 | 0.968 | 0.954 | 0.924 | 0.898 | 0.782 | 0.651 |
| **ERMN** | 1.001 | 0.998 | 1.002 | 0.992 | 0.993 | 1.003 | 0.997 |
| **FAAH2** | 0.988 | 0.974 | 0.929 | 0.916 | 0.888 | 0.780 | 0.649 |
| **FBXL16** | 0.997 | 1.017 | 1.018 | 1.019 | 1.060 | 1.054 | 1.082 |
| **FKBP5** | 0.999 | 0.996 | 0.997 | 0.999 | 0.994 | 0.978 | 0.951 |
| **FLT3** | 0.989 | 0.981 | 1.023 | 1.014 | 1.024 | 1.021 | 1.106 |
| **FOSL2** | 0.989 | 0.972 | 0.961 | 0.935 | 0.904 | 0.786 | 0.675 |
| **FPGS** | 1.010 | 1.017 | 1.035 | 1.057 | 1.078 | 1.189 | 1.420 |
| **GABARAPL1** | 0.989 | 0.981 | 0.972 | 0.954 | 0.933 | 0.861 | 0.773 |
| **GCNT4** | 1.001 | 0.998 | 1.008 | 0.998 | 1.015 | 0.990 | 1.043 |
| **HAL** | 0.994 | 0.988 | 0.981 | 0.974 | 0.965 | 0.897 | 0.815 |
| **HDAC9** | 0.999 | 1.003 | 0.996 | 0.992 | 0.982 | 0.968 | 0.936 |
| **HGS** | 0.987 | 0.971 | 0.950 | 0.925 | 0.902 | 0.793 | 0.683 |
| **HNRNPDL** | 0.984 | 0.962 | 0.941 | 0.914 | 0.876 | 0.743 | 0.634 |
| **HPCAL4** | 0.997 | 1.015 | 1.001 | 0.969 | 0.949 | 0.921 | 0.768 |
| **ID3** | 1.002 | 0.985 | 0.976 | 0.987 | 0.971 | 0.950 | 0.921 |
| **IGF2R** | 0.991 | 0.977 | 0.968 | 0.944 | 0.918 | 0.805 | 0.703 |
| **IL13RA1** | 0.987 | 0.963 | 0.962 | 0.931 | 0.888 | 0.784 | 0.669 |
| **IL18R1** | 0.990 | 0.967 | 0.943 | 0.901 | 0.898 | 0.768 | 0.642 |
| **IL1R1** | 0.989 | 0.985 | 0.957 | 0.957 | 0.932 | 0.826 | 0.750 |
| **IL1R2** | 1.004 | 1.003 | 1.007 | 1.004 | 1.008 | 0.996 | 0.987 |
| **KIDINS220** | 0.997 | 0.987 | 0.975 | 0.969 | 0.949 | 0.888 | 0.787 |
| **KLF9** | 0.996 | 0.992 | 0.983 | 0.978 | 0.957 | 0.904 | 0.829 |
| **LEF1** | 1.008 | 1.017 | 1.030 | 1.045 | 1.065 | 1.155 | 1.283 |
| **MAL** | 0.998 | 0.996 | 0.994 | 0.990 | 0.979 | 0.951 | 0.886 |
| **MEGF9** | 0.998 | 0.995 | 0.991 | 0.985 | 0.979 | 0.921 | 0.842 |
| **MFGE8** | 0.993 | 0.990 | 0.983 | 0.970 | 0.966 | 0.937 | 0.857 |
| **MGAM** | 0.988 | 0.970 | 0.952 | 0.938 | 0.909 | 0.814 | 0.694 |
| **MKNK2** | 1.005 | 1.011 | 1.021 | 1.032 | 1.044 | 1.106 | 1.199 |
| **MRFAP1** | 1.004 | 1.009 | 1.016 | 1.026 | 1.038 | 1.080 | 1.135 |
| **MSRB2** | 1.004 | 1.004 | 1.001 | 0.986 | 0.987 | 0.963 | 0.982 |
| **MTHFS** | 0.994 | 0.986 | 0.968 | 0.944 | 0.919 | 0.827 | 0.737 |
| **MYO1E** | 1.007 | 1.012 | 1.035 | 1.047 | 1.080 | 1.197 | 1.353 |

| | Downsampled Fraction | | | | | | |
|---|---|---|---|---|---|---|---|
| **Marker** | 0.9 | 0.8 | 0.7 | 0.6 | 0.5 | 0.25 | 0.1 |
| **NELL2** | 1.007 | 1.016 | 1.028 | 1.043 | 1.067 | 1.187 | 1.376 |
| **NR1D1** | 0.995 | 0.997 | 1.009 | 1.004 | 0.995 | 0.969 | 0.951 |
| **NR1D2** | 1.000 | 0.998 | 1.003 | 0.999 | 1.002 | 0.976 | 0.949 |
| **NRCAM** | 1.001 | 0.997 | 0.999 | 0.988 | 1.002 | 0.932 | 0.842 |
| **OTUD5** | 1.006 | 1.010 | 1.018 | 1.018 | 1.035 | 1.069 | 1.109 |
| **PER1** | 0.999 | 1.002 | 1.000 | 1.000 | 1.000 | 0.985 | 0.979 |
| **PER2** | 0.983 | 0.956 | 0.913 | 0.882 | 0.848 | 0.719 | 0.599 |
| **PER3** | 0.990 | 0.964 | 0.934 | 0.962 | 0.877 | 0.818 | 0.705 |
| **PFKFB2** | 1.001 | 0.992 | 0.996 | 0.970 | 0.971 | 0.905 | 0.793 |
| **PHC2** | 0.995 | 0.989 | 0.992 | 0.976 | 0.959 | 0.915 | 0.855 |
| **PPIB** | 1.006 | 1.013 | 1.021 | 1.033 | 1.044 | 1.093 | 1.158 |
| **REPS2** | 0.996 | 0.994 | 0.988 | 0.988 | 0.987 | 0.934 | 0.902 |
| **SAP30** | 0.993 | 0.989 | 0.979 | 0.967 | 0.967 | 0.927 | 0.850 |
| **SEC14L1** | 1.009 | 1.020 | 1.039 | 1.060 | 1.090 | 1.222 | 1.431 |
| **SIPA1L2** | 0.973 | 0.940 | 0.909 | 0.816 | 0.792 | 0.637 | 0.462 |
| **SMAP2** | 1.005 | 1.006 | 1.012 | 1.014 | 1.019 | 1.025 | 1.010 |
| **SPATA6** | 0.978 | 0.951 | 0.917 | 0.885 | 0.820 | 0.685 | 0.514 |
| **SYTL3** | 0.988 | 0.970 | 0.943 | 0.925 | 0.884 | 0.750 | 0.655 |
| **TBXAS1** | 1.007 | 1.020 | 1.036 | 1.056 | 1.090 | 1.235 | 1.500 |
| **TCF7** | 1.003 | 1.001 | 1.008 | 1.007 | 1.013 | 1.008 | 0.994 |
| **TMCC3** | 1.008 | 1.008 | 1.015 | 1.029 | 1.058 | 1.129 | 1.246 |
| **TPST1** | 1.009 | 1.004 | 1.005 | 1.020 | 1.035 | 1.083 | 1.069 |
| **TRABD2A** | 1.001 | 1.001 | 1.009 | 1.016 | 1.017 | 1.032 | 1.035 |
| **TRAP1** | 0.995 | 1.000 | 1.019 | 1.021 | 0.999 | 1.037 | 1.008 |
| **TSC22D3** | 1.003 | 1.006 | 1.016 | 1.023 | 1.030 | 1.085 | 1.158 |
| **UPB1** | 0.976 | 1.014 | 1.043 | 0.989 | 0.961 | 1.014 | 0.811 |
| **USP19** | 0.981 | 0.967 | 0.940 | 0.907 | 0.878 | 0.783 | 0.615 |
| **XKR8** | 1.008 | 1.019 | 1.037 | 1.043 | 1.070 | 1.163 | 1.299 |
| **ZBTB16** | 0.994 | 0.989 | 0.978 | 0.953 | 0.935 | 0.846 | 0.734 |
