## Supplementary File 3 for "TrACES of Time: Towards estimating time-of-day of bloodstain deposition by targeted RNA sequencing"

**Supplementary File 3: Library preparation QIAseq targeted RNA sequencing with UMIs**

**Step 1: Genomic DNA eliminiation**

1. Thaw Reagents and briefly centrifuge to collect at the bottom of wells
2. Prepare genomic DNA elimination mix for each sample:

| Component | Volume [µl] |
|---|---|
| **Buffer GE** | 1 |
| **Total RNA (25 ng)** | ≤ 4 |
| **Nuclease free water** | To 5 |
| **Total** | 5 |

3. Mix gently and briefly centrifuge
4. Incubate at 42°C for 5 min, then immediately place on ice for 1 min

**Step 2: Reverse Transcription**

1. Prepare Reverse Transcription mix for each sample:

| Component | Volume [µl] |
|---|---|
| **5x Buffer BC3** | 2 |
| **Control P2** | 0.5 |
| **RE3 Reverse Transcriptase Mix** | 1 |
| **RNAse free water** | 1.5 |
| **Total** | 5 |

2. Add 5 µl Reverse Transcription mix to each gDNA-eliminated sample, mix gently and briefly centrifuge
3. Incubate in thermocycler:

| Temperature | Time |
|---|---|
| 42°C | 15 min |
| 95°C | 5 min |

STOPPING POINT

➔ Store samples on ice for short term or at -15 to -30°C for longer storage

**Step 3: Molecular Barcode Assignment**

1. Prepare Barcode assignment reaction mix for each sample:

| Component | Volume [µl] |
|---|---|
| **QIAseq RNA 5x Buffer** | 4 |
| **BC Primer mix** | 2 |
| **HotStar Taq DNA Polymerase** | 0.8 |
| **DNAse-free water** | 3.2 |
| **cDNA** | 10 |
| **Total** | 20 |

2. Mix gently and briefly centrifuge
3. Place in thermal cycler with the following reaction conditions:

| Temperature | Time |
|---|---|
| 95°C | 15 min |
| 55°C | 15 min |
| 65°C | 15 min |
| 72°C | 7 min |
| 4°C | Hold |

STOPPING POINT

➔ Store samples on ice for short term or at -15 to -30°C for longer storage

**Step 4: Cleanup of BC Assignment reaction with 2 rounds of QIAseq beads purification**

1. Bring QIAseq beads to room temperature 30 min prior to use and mix well
2. Add 30 µl water to the 20 µl reaction mix (final volume: 50 µl) and mix well
3. Add 90 µl QIAseq beads (1.8 x volume) to the reaction mix and mix well
4. Incubate at room temperature for 5 min
5. Place on a 96-well magnetic rack for min. 2 minutes until solution clears
6. Discard supernatant without disturbing the beads
7. Quickspin the to settle any residual liquid to the bottom and completely remove it
8. Elute the DNA target beads in 25 µl sterile water and mix well
9. Incubate 1-5 mins at room temperature. There is no need to separate beads from eluate.
10. Add 32.5 µl (1.3 x volume) of QIAseq beads to 25µl reaction and mix well
11. Incubate at room temperature for 5 min
12. Place on a 96-well magnetic rack for min. 2 minutes until solution clears
13. Discard supernatant without disturbing the beads
14. Quickspin the tube to settle any residual liquid to the bottom and completely remove it
15. Add 200 µl freshly prepared ethanol (80%) to wells on the magnetic rack. Move the tubes from side to side between the two positions of the magnet to wash the beads, then carefully discard supernatant without disturbing the pellet.
16. Repeat step 16 for a second wash.

17. Quickspin the tube to settle any residual liquid to the bottom and completely remove it
18. Air-dry beads for 5-10 min
19. Elute DNA target beads into 12 µl sterile water, mix well, incubate at room temperature for 5 minutes then place tubes on magnetic rack until the solution is clear
20. Transfer 10 µl supernatant to a clean PCR stripe

STOPPING POINT

- ➔ Store samples on ice for short term or at -15 to -30°C for longer storage

**Step 5: 1st stage PCR**

1. Prepare 1st stage PCR mix for each sample:

| Component | Volume [µl] |
|---|---|
| **QIAseq RNA 5x Buffer** | 5 |
| **LA Primer Mix** | 2.5 |
| **RS2 Primer** | 1.5 |
| **Purified product from previous protocol** | 10 |
| **Hot Star Taq DNA Polymerase** | 1 |
| **DNAse free water** | 5 |
| **Total** | 25 |

2. Place in thermal cycler:

| Stage | Temperature | Time |
|---|---|---|
| Hold | 95°C | 15 min |
| Cycle x 12 | 95°C | 15s |
| | 60°C | 5 min |
| Hold | 4°C | Hold |

STOPPING POINT

➔ Store samples on ice for short term or at -15 to -30°C for longer storage

**Step 6: Cleanup of 1st stage PCR with 2 rounds of QIAseq beads purification**

1. Bring QIAseq beads to room temperature 30 min prior to use and mix well
2. Add 40 µl (1.6 x volume) of QIAseq beads to 25µl PCR reaction and mix well
3. Incubate at room temperature for 5 min
4. Place on a 96-well magnetic rack for min. 2 minutes until solution clears
5. Discard supernatant without disturbing the beads
6. Quickspin the tube to settle any residual liquid to the bottom and completely remove it
7. Elute the target DNA in 25 µl of water and mix well
8. Incubate 1-5mins at room temperature. There is no need to separate the beads from the eluate.
9. Add 40 µl (1.6 x volume) of QIAseq beads to the 25 µl reaction and mix well
10. Incubate at room temperature for 5 minutes.
11. Place on a 96-well magnetic rack for min. 2 minutes until solution clears
12. Discard supernatant without disturbing the beads
13. Add 200 µl freshly prepared ethanol (80%) to wells on the magnetic rack. Move the tubes from side to side between the two positions of the magnet to wash the beads, then carefully discard supernatant without disturbing the pellet.
14. Repeat step 16 for a second wash.
15. Quickspin the tube to settle any residual liquid to the bottom and completely remove it

16. Air-dry beads for for 5-10 min
17. Elute DNA target beads into 27 µl sterile water, mix well by pipetting, incubate at room temperature for 5 minutes then place tubes on magnetic rack until the solution is clear
18. Transfer 25 µl supernatant to a clean PCR stripe

STOPPING POINT

- ➔ Store samples on ice for short term or at -15 to -30°C for longer storage

**Step 7: 2nd stage Universal Index PCR**

1. Prepare 2nd stage PCR mix for each sample in barcode array plate.

| Component | Volume [µl] |
|---|---|
| **QIAseq RNA 5x Buffer** | 10 |
| **FS-trP1** | - |
| **RS-ID#** | - |
| **Purified product from previous protocol** | 25 |
| **Hot Star Taq DNA Polymerase** | 2 |
| **DNAse free water** | 13 |
| **Total** | 50 |

#ID1-ID96 (adapted from Ion Xpress 1- 96)

2. Thermal cycler program:

| Stage | Temperature | Time |
|---|---|---|
| Hold | 95°C | 15 min |
| Cycle x 20 | 95°C | 15s |
| | 60°C | 2 min |
| Hold | 4°C | Hold |

STOPPING POINT

➔ Store samples on ice for short term or at -15 to -30°C for longer storage

**Step 8: Cleanup of universal PCR with 1 round of QIAseq beads purification**

1. Bring QIAseq beads to room temperature 30 min prior to use and mix well
2. Add 55 µl (1.1 x volume) of QIAseq beads to 50µl PCR reaction and mix well
3. Incubate at room temperature for 5 min
4. Place on a 96-well magnetic rack for min. 2 minutes until solution clears
5. Discard supernatant without disturbing the beads
6. Quickspin the tube to settle any residual liquid to the bottom and completely remove it
7. Add 200 µl freshly prepared ethanol (80%) to wells on the magnetic rack. Move the tubes from side to side between the two positions of the magnet to wash the beads, then carefully discard supernatant without disturbing the pellet.
8. Repeat step 16 for a second wash.
9. Quickspin the tube to settle any residual liquid to the bottom and completely remove it
10. Air-dry beads for for 5-10 min
11. Elute DNA target beads into 27 µl sterile water, mix well by pipetting, and incubate at RT for 15 minutes
12. then place tubes on magnetic reack until the solution is clear
13. Transfer 25 µl supernatant to a clearn PCR stripe

STOPPING POINT

- ➔ Store samples on ice for short term or at -15 to -30°C for longer storage.

**Step 9: Library Quantification**

1. Prepare a serial dilution for Ion Torrent DNA standard:

| Standard | DNA Standard | Nuclease-free water |
|---|---|---|
| 1 | 5 µl undiluted | 45 µl |
| 2 | 5 µl Std. 1 | 45 µl |
| 3 | 5 µl Std. 2 | 45 µl |
| 4 | 5 µl Std. 3 | 45 µl |
| 5 | 5 µl Std. 4 | 45 µl |

2. Prepare dilutions for each sample library:

| Dilution | Library | Nuclease-free water |
|---|---|---|
| Starting Dilution (1:20) | 2 µl undiluted | 38 µl |
| Dilution 1 (1:2,000) | 2 µl of 1:20 | 198 µl |
| Dilution 2 (1:20,000) | 5 µl of 1:2.000 | 45 µl |

3. Prepare a PCR mastermix

| Component | Volume [µl] |
|---|---:|
| **RNAse/DNAse free water** | 2.75 |
| **SYBR Green Mastermix** | 6.25 |
| **Primermix** | 0.5 |
| **Total** | 22 |

4. Add 3 µl of sample per reaction well
5. Tightly seal the reaction plate
6. Centrifuge for 1 min at 1000 x g to remove any bubbles
7. Start qPCR run with the following cycling conditions

| Stage | Temperature | Time |
|---|---|---|
| Hold | 95°C | 10 min |
| Cycle x 30 | 95°C | 15 s |
| | 60°C | 30 s |
| | 72°C | 2 min* |

*collect fluorescence data

8. Calculate library concentration using the "GeneRead Library Quant Kit" Data Analysis File (available from: https://www.qiagen.com/us/products/human-id-and-forensics/nextgeneration-sequencing/qiaseq-library-quant-system (last accessed: 2025-01-03))

**Step 10: Library Dilution**

1. Dilute each individual library to 50 pM in nuclease-free water
